# tACS induces state-dependent shifts in inhibitory phase preference consistent with phase precession

**DOI:** 10.64898/2025.11.28.691215

**Authors:** Zhou Fang, Sanne ten Oever, Alexander Sack, Inge Leunissen

## Abstract

Phase-dependent modulation of motor inhibition suggests that oscillatory timing plays a key role in inhibitory control. Here we show that rhythmic brain stimulation can selectively alter this timing in a manner consistent with phase precession. Participants performed an anticipatory stop-signal task while receiving 20 Hz transcranial alternating current stimulation (tACS) over the pre-supplementary motor area, with stop signals presented at defined phases of the stimulation waveform. An additional control experiment with no tACS was conducted, in which the stop signal phase was derived post-hoc from the ongoing EEG. In both tACS and control sessions, stopping efficacy varied sinusoidally with beta phase, confirming that inhibitory control is intrinsically phase-dependent. During tACS, however, the phase associated with the strongest inhibition advanced progressively across the session by about 180°, and this effect was specific to failed-stop trials. No comparable shift occurred without stimulation. These findings indicate that rhythmic drive can bias the temporal phase at which inhibition is most effective when the brain is in an error-prone state, revealing a state-dependent, precession-like reorganization of inhibitory timing. We propose that tACS interacts with error-related plasticity to promote gradual phase advancement, placing the system within spike-timing-dependent learning windows, whereas intrinsic fluctuations alone appear insufficient to facilitate cumulative precession. More broadly, these results suggest that phase precession may represent a general coding principle extending beyond hippocampal circuits, enabling the motor system to adaptively adjust inhibitory control through oscillatory timing mechanisms.

**Highlights:**

- Beta tACS over preSMA induces shifts in phase preference during motor inhibition
- Precession-like phase shifts emerge selectively during failed but not successful stops
- Phase precession was absent in a control experiment without tACS
- tACS might provide a method to probe phase precession mechanisms in cognition

## Introduction

Neural oscillations are fundamental to brain function, providing a temporal framework that regulates the timing of neuronal activity and thereby supports synaptic plasticity, information transfer, and large-scale integration across cortical and subcortical networks (Buzsaki and Draguhn, 2004; Beste et al., 2023). One striking oscillatory phenomenon is phase precession, a coding mechanism in which neural spikes shift progressively earlier relative to the phase of an ongoing oscillation across successive cycles. This phenomenon was first described in the hippocampus of rodents during spatial navigation (O’Keefe and Recce, 1993) and has since been observed in regions such as the ventral striatum (van der Meer and Redish, 2011) and frontal cortex (Jones and Wilson, 2005), suggesting that it represents a general computational principle extending beyond the hippocampal system. By advancing spike timing across successive oscillatory cycles, neurons can represent behavioral sequences that unfold over longer timescales within the faster temporal structure of ongoing rhythms (Lisman and Jensen, 2013; Widloski and Fiete, 2014). This temporal compression is thought to “bind” distributed neural events into coherent, rapidly accessible representations, supporting associative learning and flexible behavior (Lisman and Jensen, 2013; Widloski and Fiete, 2014; Buzsaki and Tingley, 2018; Schlesiger et al., 2015). Recent invasive recordings have further demonstrated phase precession in the human hippocampus (Qasim et al., 2021; Reddy et al., 2021), underscoring its relevance for human neural coding.

A critical next step is to determine whether phase precession in humans can be induced or promoted experimentally. Establishing causal control over this phenomenon would be important not only for testing its proposed roles in sequence learning, temporal binding, and predictive coding, but also for exploring how this mechanism might be harnessed to modulate cognition and behavior. Transcranial alternating current stimulation (tACS) provides a promising approach. By applying low-intensity alternating currents at specific frequencies, tACS can bias ongoing rhythms, promote phase alignment of neuronal spiking time (Ozen et al., 2010; Johnson et al., 2020; Krause et al., 2019), and modulate brain oscillations through a process known as entrainment (Thut et al., 2011; Herrmann et al., 2016). At the cellular level, alternating currents have been shown to advance spike phase relative to an imposed oscillation, providing a mechanistic precedent for externally driven precession (Magee, 2001; Narayanan and Johnston, 2008). While most human tACS studies have demonstrated stable phase alignment rather than progressive phase shifts, recent work has shown that rhythmic stimulation can induce phase precession in the motor cortex (Wischnewski et al., 2024). This raises the question of whether tACS-induced phase precession generalizes beyond simple motor cortex activity and whether its expression is directly linked to cognitive function and behavior. Building on these findings, the present study investigates whether tACS can enable phase precession-like dynamics during inhibitory control. Specifically, we tested whether rhythmic stimulation over the pre-supplementary motor area (preSMA) could induce systematic phase advancement during motor inhibition, a process in which stopping efficiency has been shown to depend on the beta oscillatory phase (Fang et al., 2024). We propose that tACS provides a rhythmic scaffold that aligns neural activity in time, allowing intrinsic excitatory-inhibitory interactions to gradually advance the phase of optimal motor inhibition within this externally synchronized rhythm. Such temporal compression of neural activity has been proposed to facilitate spike-timing-dependent plasticity and adaptive coding in animal models (Dan and Poo, 2006; Lengyel et al., 2005). Accordingly, we hypothesized that if tACS promotes phase precession-like dynamics, inhibitory performance should exhibit a progressive shift over time in the optimal stimulation phase at which the stop-signal arrives. In contrast, this shift should be absent without external rhythmic drive in a control experiment, indicating that precession reflects an interaction between stimulation-induced synchronization and endogenous network adaptation.

## Materials and Methods

### Overview

This study comprised two experiments. In Experiment 1, we reanalyzed the dataset reported in Fang et al. (2024) to address a distinct question regarding tACS-induced phase precession. The experimental design has been described in detail previously (Fang et al., 2024) and is summarized here. In Experiment 2, an independent cohort performed the same stop-signal task but without active tACS, providing a control condition. Common procedures and key differences between the two experiments are outlined below.

### Participants

In Experiment 1, twenty-nine healthy, right-handed adults (13 males, 19 females; mean age = 25.3 years, range 18-35) participated. Handedness was confirmed using the Edinburgh Handedness Inventory (mean laterality quotient = 96.3, range 70-100; (Oldfield, 1971)). In Experiment 2, eighteen healthy, right-handed adults (6 males, 12 females; mean age = 24.0 years, range 18-30, mean laterality quotient = 89.4, range 65-100) participated. All participants had no contraindications to non-invasive brain stimulation (Bikson et al., 2009; Woods et al., 2016), and provided written informed consent. The protocol was approved by the Ethics Review Committee Psychology and Neuroscience of Maastricht University, the Netherlands (ERCPN Approval code: OZL_204_04_02_2019).

### Task and stimulation protocol

In both experiments, participants performed an anticipatory version of a stop signal task (Slater-Hammel, 1960; Coxon et al., 2006; Leunissen et al., 2017). As shown in the Figure 1, the trial starts with a moving indicator filling up an empty bar (1000 ms) with a consistent speed. Participants were asked to stop the indicator as close to a fixed target line (800 ms) as possible by pinching a force transducer (OMEGA Engineering, Norwalk, CT, USA) using their right thumb and index finger. Immediate feedback indicated absolute error: green, yellow, orange, or red for ≤20, ≤40, ≤60, or *>*60 ms, respectively. Response was triggered when their force exceeds a threshold which was determined from individual maximal voluntary force (MVF). MVF was defined as the highest value obtained across three 5 s maximal pinches. The threshold was initially set to 25% MVF and could be reduced in 5% increments for comfort (mean 23.3%, range 10-25%). In 30% of the trials, the bar stopped before reaching the target, signaling participants to withhold their response. Failed stop trials (SF) were those where force exceeded the response thresholds, while successful stop trials (SS) involved forces below this threshold.

**Figure 1.**
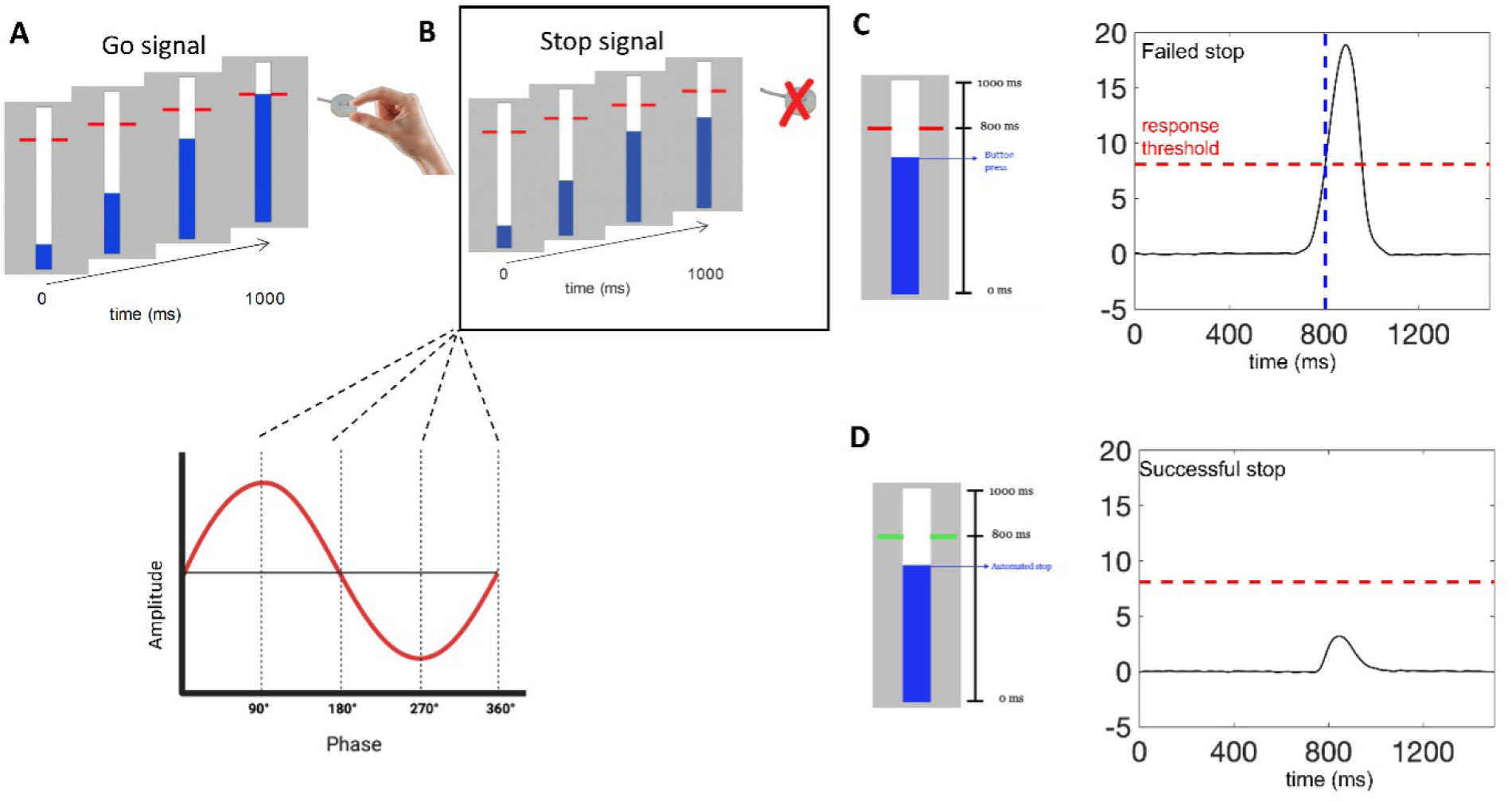
Anticipatory stop-signal task and trial classification. (**A**) In the go trials, participants squeezed a force sensor as a bar reached the target. (**B**) In stop trials (30% of the total trials), the bar stopped early, requiring inhibition. In Experiment 1, the stop signals were presented equiprobably at four phase angles (90◦, 180◦, 270◦, 360◦) relative to the tACS; in Experiment 2, the instantaneous phase at the stop signals was estimated post hoc and assigned to the same four phase bins. (**C**) Stop-failed (SF) were defined as responses exceeding the response threshold. The target line turned red to indicate failed inhibition. (**D**) Stop-successful (SS) were those in which force remained below the threshold; the target line turned green to indicate successful inhibition.

In Experiment 1, the session comprised 240 stop trials and 480 go trials presented in pseudorandom order. Stop-signal delays (SSD) occurred approximately 110, 150, 190, 230, and 270 ms prior to the target line and shifted up to ±25 ms to match the intended tACS phase. Inter-trial intervals jittered between 2.7 and 3.7 s. tACS (DC STIMULATOR PLUS, NeuroConn, GmbH, Ilmenau, Germany) was placed using a 1×4 electrode configuration over the preSMA, with the central electrode located at FCz and four surrounding electrodes at F1, F2, C1, and C2 based on the 10-20 system. The electrodes were embedded in gel-filled cup that was made out of plastic cylinders (,02 cm at top, 2.5 cm at bottom) and mounted in an EEG cap (EASYCAP GmbH, Germany). The stop signals were presented at 90◦ (peak), 180◦, 270◦, and 360◦ relative to the tACS waveform at 20 Hz using a phase-locked stimulus presentation system (ten Oever et al., 2016), as shown in FIgure.1 (B).

Experiment 2 involved 101 stop trials and 308 go trials, presented in 3 blocks of 136 trials. We used a staircase tracking algorithm that adjusts the SSD in 16.667 ms increments based on the participant’s performance to maintain a 50% inhibition accuracy in stopping responses on stop trials. During the experiment, tACS electrodes were positioned as in Experiment 1. Stimulation was delivered only during the first 10 seconds of each task block, leaving the remainder 10 min tACS-free. For the purpose of estimating the stop-signal phase relative to the ongoing oscillatory phase, EEG was recorded using a BrainAmp system (Brain Products GmbH, Germany) at a sampling rate of 5000 Hz from 32 Ag/AgCl electrodes according to the 10-20 system (channels included F7, FC5, C3, Fz, FC1, Cz, FC2, C4, FC6, F8, T7; T8 as implicit reference; AFz as ground). In addition, four electro-oculographic (EOG) electrodes (UVEOG, LVEOG, RHEOG, and LHEOG) were placed to monitor eye movements. Impedances were maintained below 10 kΩ and signals were sampled at 5000 Hz. Processing performed in MATLAB R2022a using FieldTrip (Oostenveld et al., 2011) comprised (1) re-referencing each channel to T7; (2) demeaning and detrending; (3) filtering with a 1-300 Hz band pass filter and a 50 Hz FIR notch filter; (4) regressing out eye movement noise by an EEGlab plugin, AAR (order = 3, *λ* = 0.9999, *σ* = 0.01, precision = 50). The phase at the moment of the stop signal was determined post-hoc based on the Fz channel: it was first segmented from –0.5 to 0 s relative to the stop signal, and a one-pass FIR bandpass filter (18-22 Hz) was applied. The edge of 32 data points preceding the stop signal was removed to mitigate boundary artifacts introduced by filtering. An autoregressive (AR) model (order 30) was then fitted to predict 64 data points forward. The Hilbert transform was applied to the segment, estimating the phase at the instant corresponding to the original endpoint. Estimated phases were categorized into four bins: 90*^◦^* (≥ 45*^◦^* to *<* 135*^◦^*), 180*^◦^* (≥ 135*^◦^* to *<* 225*^◦^*), 270*^◦^* (≥ 225*^◦^* to *<* 315*^◦^*), and 360*^◦^* (≥ 315*^◦^* to *<* 360*^◦^* or ≥ 0*^◦^* to *<* 45*^◦^*).

### Force metrics

The force signal was acquired via a digital-to-analog converter (DAC) from National Instruments (Austin, TX, USA) and recorded in LabChart with a sampling rate of 1000 Hz. Data were saved on a trial-by-trial basis from the onset of the fill indicator to 1700 ms.

Behavioral force data were analyzed using MATLAB R2022a (MathWorks, Natick, MA, USA). For both Experiment 1 and 2, raw force signals recorded on a trial-by-trial basis were first converted from volts to newtons. A fifth-order 20 Hz low-pass Butterworth filter was applied to remove high-frequency noise. Baseline correction was performed by subtracting the mean force measured between −650 ms and −400 ms relative to target onset for each trial.

For all stop trials, the peak force, i.e. the maximum force within the trial, was extracted and normalized to the participant’s MVF, yielding a percentage measure. Trials with force values exceeding ±2.5 standard deviations from the participant’s mean were considered outliers and removed from further analysis.

### Data analysis

Phase dependency and phase precession of motor inhibition were both quantified from normalized peak force values across the four stop-signal phase bins (90*^◦^*, 180*^◦^*, 270*^◦^*, 360*^◦^*). The phase dependency analysis assessed how stopping efficacy varied across these phases, whereas phase precession was evaluated using the same procedure but within a sliding time window across the experimental session.

### Phase dependency analysis

For each experiment, normalized peak force values were compared across the four phase bins. In Experiment 1 (SS), data violated the assumption of normality (Shapiro-Wilk test, *p <* 0.05); therefore, a Friedman test (non-parametric test for repeated measures) was used. For Experiment 1 (SF) and Experiment 2, a repeated-measures ANOVA (rm-ANOVA) was applied. Post hoc tests examined pairwise differences between phase bins. To characterize the sinusoidal modulation of inhibition, normalized peak force values were first averaged across participants within each phase bin, and a sine wave was fitted to the group means. The goodness-of-fit (*R*^2^) was compared against a permutation-based null distribution obtained by shuffling phase labels at the individual level to assess the reliability of the modulation. The preferred phase was defined as the trough of the fitted sine wave (reflecting the phase of strongest inhibition), and the weight of the preferred phase was taken as the amplitude of the sine wave, indicating the magnitude of modulation.

### Phase precession analysis

Phase precession was assessed by re-computing the preferred phase and weight within a sliding window across the experiment. In each window, a sine wave was fitted to normalized peak force values averaged across participants for the four phase bins, and the trough and amplitude were extracted as the preferred phase and weight, respectively. In Experiment 1, each window included at least 80 stop-signal trials (covering four phase bins, five SSDs, and two conditions: SS and SF). A final window size of 96 trials (40% of total; 16 steps of 9 trials) was used. In Experiment 2, the staircase procedure concentrated SSDs near the center of the distribution; to maintain comparability, a minimum window of 48 trials was selected, corresponding to a window size of 50 trials (≈49.5% of total; 8 steps of 6 trials). Analyses were conducted separately for SS and SF trials.

Finally, linear-circular correlations between fitted preferred phases and trial progression were computed at the group level for SS and SF trials in both experiments. The statistical significance of the phase precession effect was determined by a nonparametric permutation test: phase labels were shuffled 10,000 times at the individual trial level to generate a null distribution, and *p*-values were obtained as the proportion of permutations exceeding the observed correlation coefficient. Additional analyses using different window sizes are reported in the Supplementary Information.

## Results

### Experiment 1 (tACS)

We first tested whether the phase of tACS at stop-signal onset influences inhibitory performance. Figure 2 (A) and (B) show the averaged SS and SF normalized peak force across stop signal phases with the best-fitting sinusoidal curve. We found a significant modulation of normalized peak force by stimulation phase for SS trials (*χ*^2^(3*, N* = 28) = 31.757, *p <* 0.001, Shapiro–Wilk test *p <* 0.05) and SF trials (*F* (3, 81) = 3.016, *p* = 0.050). Post-hoc Wilcoxon signed-rank tests confirmed significant pairwise differences between phases, indicating phase-dependent force modulation under tACS (see Table S1). Specifically, normalized peak force in SS trials was significantly lower at 90*^◦^* compared to 180*^◦^* (*p* = 0.009), 270*^◦^* (*p <* 0.001), and 360*^◦^* (*p <* 0.001). In addition, force at 180*^◦^* was lower than at 270*^◦^* (*p* = 0.036), while 270*^◦^* showed higher force than 360*^◦^* (*p* = 0.031). In SF trials, the force at 90*^◦^* was significantly higher than at 180*^◦^* (*p* = 0.001). Sinusoidal fits captured the phase-dependent modulation robustly (SS: *R*^2^ = 1.00, *p <* 0.001; SF: *R*^2^ = 0.99, *p <* 0.001), indicating significant modulation relative to a permutation-based null distribution. The trough (preferred) phases occurred at 67.3*^◦^* for SS and 132.8*^◦^* for SF trials, reflecting the phases of maximal inhibition.

**Figure 2.**
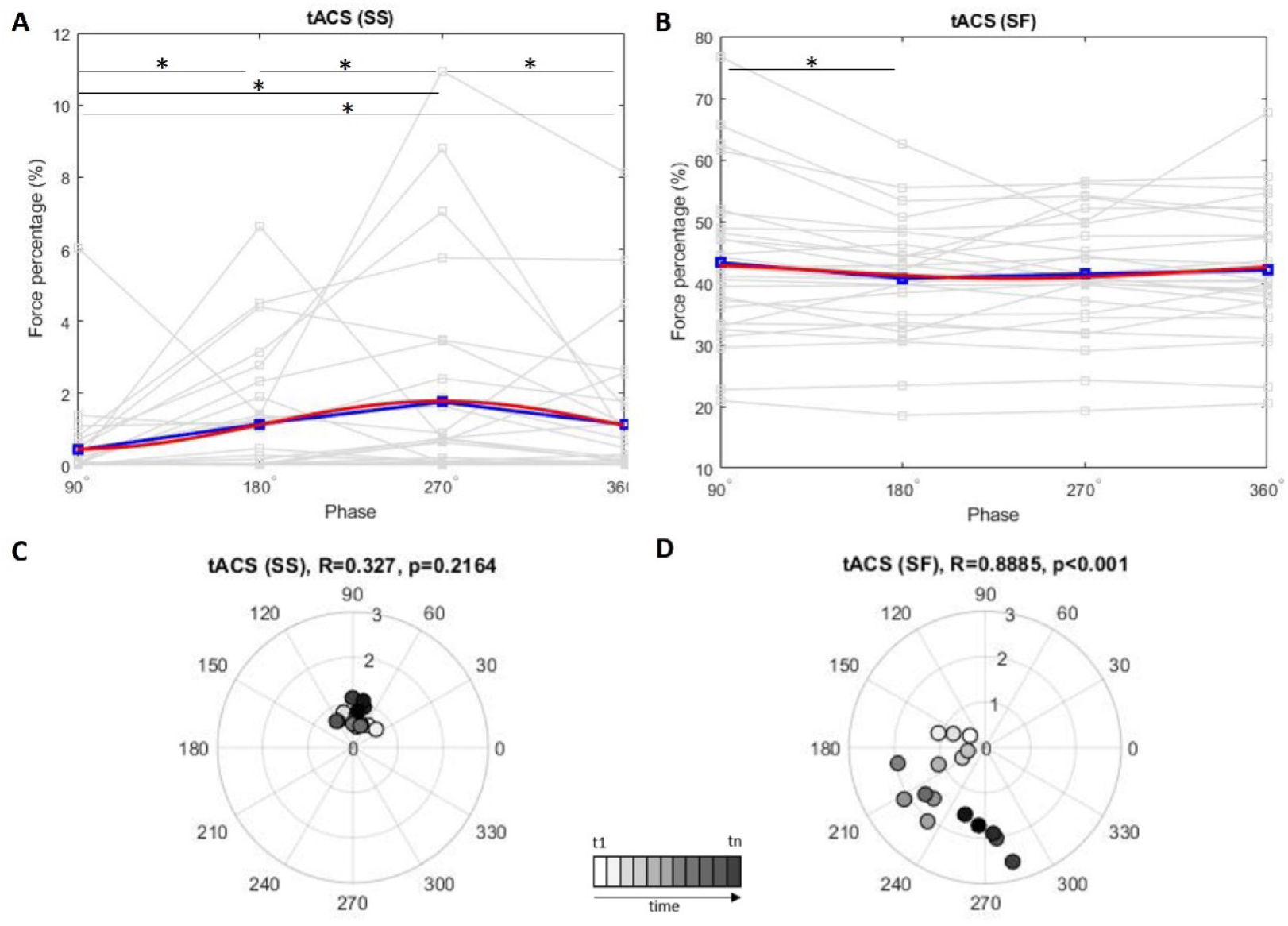
Experiment 1. (**A**)–(**B**), plots of peak force normalized with respect to the maximal voluntary force across tACS beta phases for stop success (SS) and stop failure (SF) trials. Gray lines represent individual data points. The blue lines indicate the average data, while the red curve shows the best-fitting sinusoidal pattern for each condition. Less force refers to greater inhibition. (**C**)–(**D**), polar plots of phase shifting of inhibition over time. Angular position represents preferred phase of inhibition, and radius represents the weight. A significant shift is observed in the SF group (∼120*^◦^* to ∼300*^◦^*, *R* = 0.889, *p <* 0.001), but not the SS (*R* = 0.327, *p* = 0.216).

To assess temporal changes in phase-dependent inhibition, we performed a sliding-window analysis (96 trials, 15 steps). Normalized force values were averaged across trials within each stimulation-phase bin and then across participants to determine the phase associated with the lowest force on stop trials, indexing the point of strongest inhibition. In failed stop (SF) trials, this optimal phase shifted progressively forward over the session (from approximately 120*^◦^* to 300*^◦^*), suggesting a gradual drift in the phase associated with maximal inhibition (Linear–circular correlations: *R* = 0.887, *p <* 0.001). This was not evident in SS (tACS) trials (*R* = 0.327, *p* = 0.216) (Figure 2 (C)–(D)). This pattern is held under multiple analysis windows (Figure S1). A permutation test with 10,000 shuffled phase labels at the individual trial level confirmed the significance of this effect in SF trials only (*p <* 0.001) (Figure S3).

These results suggest that under tACS, the beta phase associated with the strongest inhibitory drive shifted forward across failed inhibition trials by time, implying that error-prone states might be accompanied by preferred phase recalibration. The absence of precession in SS indicates that low-error states maintain the spike-phase relationship supporting inhibitory control.

### Experiment 2 (sham-tACS)

In the no-tACS experiment, we also first assessed whether inhibitory performance depends on the stop-signal phase. As shown in Figure 3 (A)-(B), force output during SS trials exhibited a significant phase-dependent modulation (rm-ANOVA: *F* (3, 51) = 3.170, *p* = 0.05). Post hoc comparisons revealed significantly lower force in the 90*^◦^* bin compared to both 180*^◦^* (*p* = 0.038) and 270*^◦^* (*p* = 0.020, Table S2), suggesting enhanced inhibition when the stop signal coincided at 90*^◦^*. SF trials did not show significant phase-dependent modulation (*F* (3, 51) = 0.210, *p* = 0.889). The best-fitting sinusoidal functions showed significant phase-dependent modulation (SS: *R*^2^ = 1.00, *p <* 0.001; SF: *R*^2^ = 1.00, *p <* 0.001) relative to a permutation-based null distribution. Preferred trough phases were 82.5*^◦^* for SS and 357.8*^◦^* for SF trials, indicating maximal inhibition.

**Figure 3.**
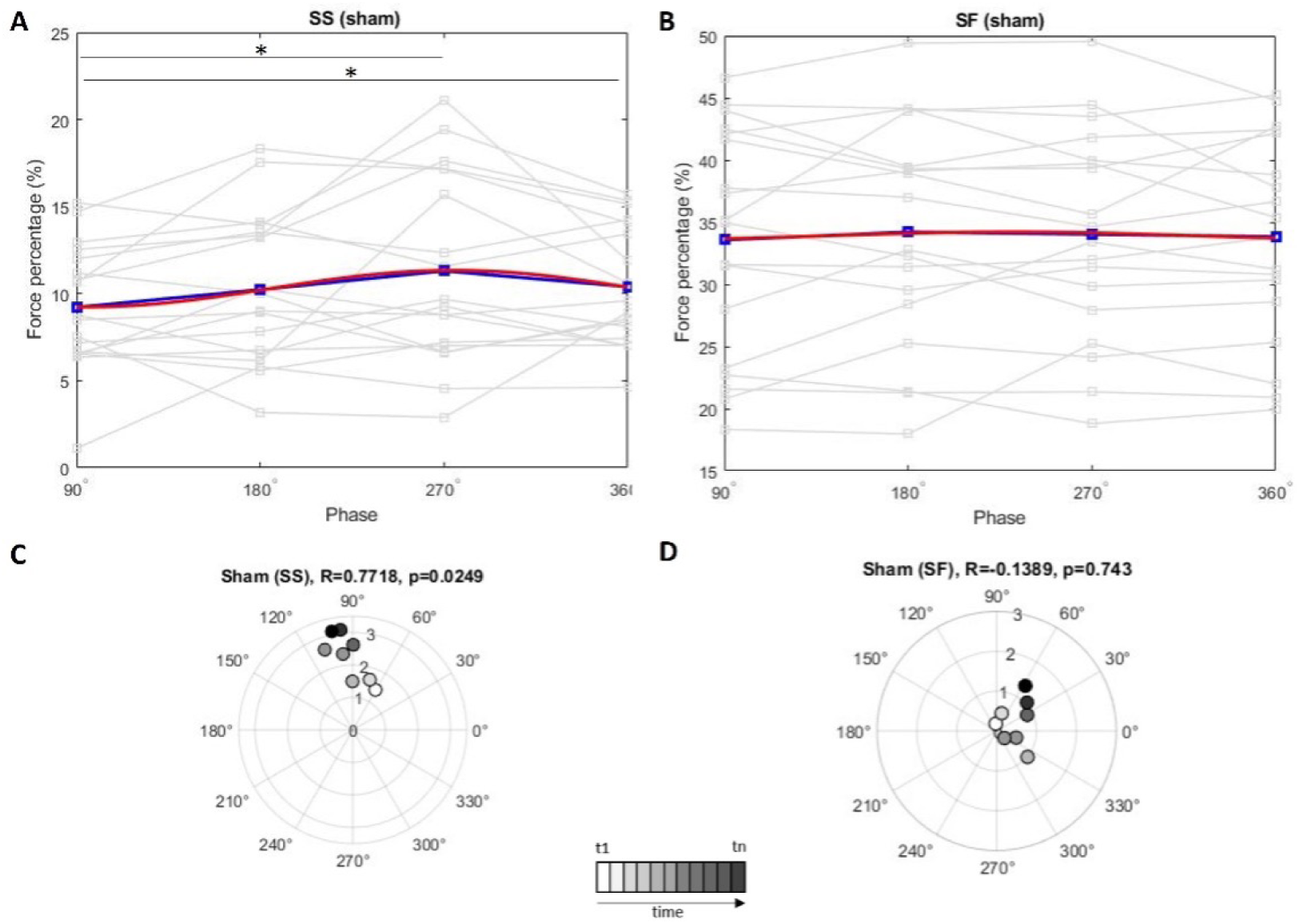
Experiment 2. (**A**)–(**B**), plots of normalized peak force across Fz-EEG beta phases for stop success (SS) and stop failure (SF) trials. Gray lines represent individual data points. The blue lines indicate the average data, while the red curve shows the best-fitting sinusoidal pattern for each condition. (**C**)–(**D**), polar plots of phase shifting of inhibition over time.

Results in Experiment 2 show successful stopping remains phase dependent with a stable optimum at ∼ 90*^◦^*. But there is no robust evidence for phase precession without stimulation. Taken together, the phase dependence of inhibition is a common property of the SS condition across experiments. The optimal beta oscillatory timing for successful stopping is stable irrespective of stimulation. SF trials showed moderate phase dependency under tACS, while no effect in the sham condition. Nevertheless, the relative phase of greatest inhibitory engagement occurred at a distinct angle compared to SS trials in both experiments, suggesting differential network recruitment between successful and failed inhibition (Cai et al., 2014). Interestingly, SF trials in the tACS condition (Experiment 1) showed a systematic advance of the optimal phase and survived rigorous permutation testing, implying that the emergence of phase precession is specific to tACS and selectively during trials where inhibitory control fails.

## Discussion

The present study tested whether externally applied rhythmic stimulation can shift the optimal oscillatory phase at which inhibitory control is strongest. Our previous work showed that stopping efficiency depends on the beta phase at which the stop-signal is presented (Fang et al., 2024)). Consistent with this, we observed a robust phase dependency of inhibitory performance in both the tACS and control experiments, confirming that phase-dependent inhibition is an intrinsic property of the motor system. Importantly, during tACS, the phase associated with the strongest inhibition advanced progressively over time ∼ 180*^◦^*, and this shift was specific to stop-failed (SF) trials. No comparable effect was observed without stimulation. This pattern suggests that external rhythmic stimulation can selectively induce the precession of phase at which inhibitory control is most effective during unsuccessful inhibition, gradually altering the coupling between oscillatory activity and behavioral performance. Although our measures do not capture neuronal spiking directly, these behavioral phase shifts are consistent with a precision-like adjustment in the functional timing of inhibitory control.

### External rhythmic stimulation alters phase preference of inhibitory control

Our work extends previous findings that beta-band stimulation can modulate motor system function at multiple levels. tACS has been shown to influence motor excitability (Wischnewski et al., 2019; Schilberg et al., 2018), alter motor performance (Pogosyan et al., 2009; Van Hoornweder et al., 2025), and affect inhibitory control (Leunissen et al., 2022; Zhu et al., 2025). Here we go beyond these observations by showing that rhythmic stimulation not only entrains oscillatory activity but can also bias the temporal phase at which inhibitory control is most effective when the brain is at an error-prone state, revealing a state-dependent modulation of oscillatory timing.

In line with Wischnewski et al. (2024), our findings support the notion that tACS can induce precession-like phase shifts, not only at rest but also during active motor inhibition, with stronger effects observed during failed inhibition trials. A possible mechanistic account is that rhythmic stimulation may create network conditions conducive to precession-like dynamics, rather than directly imposing phase shifts. In hippocampal models, phase precession has been attributed to asymmetric excitation-inhibition interactions, where excitatory ramping activity gradually outpaces inhibitory feedback, causing neurons to fire at progressively earlier oscillatory phases (Mehta et al., 2002). By synchronizing membrane potential fluctuations across populations, tACS could enhance such asymmetries, effectively amplifying small phase advances that accumulate over time. In this view, rhythmic stimulation provides a temporal scaffold within which intrinsic network dynamics can reorganize, giving rise to precession-like phase advancement without direct control of spike timing.

The predominance of precession-like phase advancement during failed stops suggests that tACS interacts most effectively with networks in an adaptive, plastic state. Failed inhibition is accompanied by dopaminergic learning signals and transient reorganization of beta activity (Schroll et al., 2018), reflecting ongoing error-driven updating of inhibitory control. During this phase, excitatory-inhibitory balance and oscillatory coupling likely fluctuate, placing neuronal populations within spike-timing-dependent plasticity (STDP) windows in which small timing shifts can alter synaptic strength (Dan and Poo, 2006; Widloski and Fiete, 2014). tACS, by providing a weak but consistent rhythmic input, may bias these timing relationships, promoting earlier activation on subsequent cycles and thereby amplifying endogenous phase advancement.

At the network level, failed inhibition also entails broader recruitment of fronto-basal ganglia and salience circuits (Samsonovich and McNaughton, 1997; Cai et al., 2014; Limongi and Pérez, 2017; Isherwood et al., 2025). This broader and more variable network configuration introduces phase heterogeneity and reduces global phase stability, making the system more susceptible to external modulation. In such a regime, a weak periodic drive can steer the relative timing among coupled oscillators, allowing cumulative phase advances to emerge over time. By contrast, successful stopping probably engages a more phase-stable, beta-synchronized circuit that resists such perturbation. This trial-specific susceptibility mirrors the state-dependent effects frequently reported in tACS studies (Krause and Cohen Kadosh, 2014; Alagapan et al., 2016), where stimulation interacts most strongly with networks that are already dynamically engaged in adaptation or learning. Together, these properties may help to explain why precession-like dynamics were observed specifically during failed stops, when both local plasticity and large-scale network variability could create conditions favorable for tACS-induced timing shifts.

### Broader theoretical implications

Our findings contribute to a growing theoretical framework in which phase precession serves as a general temporal coding mechanism in the brain. In the hippocampus, phase precession is thought to compress temporally extended events within individual oscillatory cycles, thereby supporting sequence learning and predictive coding (Widloski and Fiete, 2014). By mapping sequences of experience onto the phase of an ongoing oscillation, neural systems can link distributed events across time and space into coherent representations.

A similar principle may apply to motor inhibition. During failed stops, precession-like shifts may serve to temporally compress error-related neural dynamics, enabling the system to rapidly integrate these events into adaptive control policies. Such temporal compression could enhance the salience of behavioral errors, facilitating reinforcement-based updating of inhibitory strategies. More broadly, this framework supports the idea that phase precession is not restricted to hippocampal navigation or memory but may represent a domain-general coding principle for aligning temporally distributed neural events. From this perspective, the phase advancement observed during tACS may reflect a fundamental property of neural computation, i.e., the ability to use oscillatory phase as a temporal reference frame for linking sequential events across multiple timescales. As a result, precession enables more efficient temporal alignment of error-related neural events, potentially supporting precise action-outcome encoding and rapid adaptation for following errors. This coding strategy could underlie diverse cognitive operations, from motor adaptation and executive control to reinforcement learning and predictive processing, suggesting that oscillatory phase alignment may be a universal mechanism for temporal organization, adaptation, and plasticity in the brain.

### Limitations and future directions

In a single stimulation session, the effects of tACS on inhibitory timing are likely transient, reflecting short-term adjustments in network synchrony and excitability. With repeated exposure, however, such phase-specific modulation could begin to engage longer-term plasticity mechanisms, leading to more stable adaptations in inhibitory control. The link between precession-like dynamics and spike-timing-dependent plasticity (STDP) supports this possibility (Lengyel et al., 2005; Wischnewski et al., 2024): by aligning neural activity within potentiation windows, rhythmic stimulation may gradually reinforce phase relationships that enhance inhibitory performance. In this way, tACS could not only modulate ongoing oscillations but also shape the temporal organization of neural activity, potentially improving network efficiency over time. These considerations imply that tACS might serve to strengthen the individual capacity of error-based learning.

However, several limitations warrant caution. Inference about phase precession in the present study is indirect, relying on behavioral performance and modeled oscillatory phase rather than direct recordings of spiking activity. Consequently, the observed phase shifts should not be interpreted as evidence of neuronal precession but rather as macroscopic signatures of gradual temporal reorganization under rhythmic drive. Future studies combining tACS with invasive electrophysiology or high-resolution source imaging will be crucial to determine whether similar progressive phase shifts occur at the neuronal ensemble level.

An additional possibility is that the observed phase advancement arises from changes in coupling strength between the exogenous tACS field and endogenous oscillations rather than from intrinsic timing adjustments. According to coupled-oscillator models such as the Kuramoto framework (Kuramoto, 1984), dynamic fluctuations in coupling can shift the phase relationship between two oscillators without altering intrinsic spike timing. Although such a mechanism could, in principle, explain gradual phase drift, the outcome-specific pattern observed here argues against a purely coupling-based explanation: during successful-stop trials, the tACS-EEG beta phase relationship remained stable.

Future work combining rhythmic stimulation with computational modeling, MEG/EEG source reconstruction, and invasive recordings will be essential to distinguish genuine intrinsic precession from changes in network-level coupling. These approaches will also help determine whether repeated induction of precession-like shifts can produce lasting improvements in inhibitory control and behavioral flexibility, offering a pathway toward rhythm-based modulation of adaptive cognition.

## Conclusion

Taken together, our results show that neuronal phase preference under tACS can shift over time during goal-directed inhibition, and that this effect is contingent on behavioral outcome. These findings suggest that phase precession is not only a biophysical phenomenon but also one shaped by cognitive demands and capable of interacting with plasticity mechanisms. More broadly, our work provides a methodological framework for probing phase dynamics in motor control and highlights the potential of outcome-contingent stimulation paradigms to harness precession as a mechanism for adaptive neuromodulation.

## Supporting information

Latex project

## Supplementary Information

### Supplementary 1 – Post hoc Comparisons of Force Modulation by Beta Phase

**Table S1.**
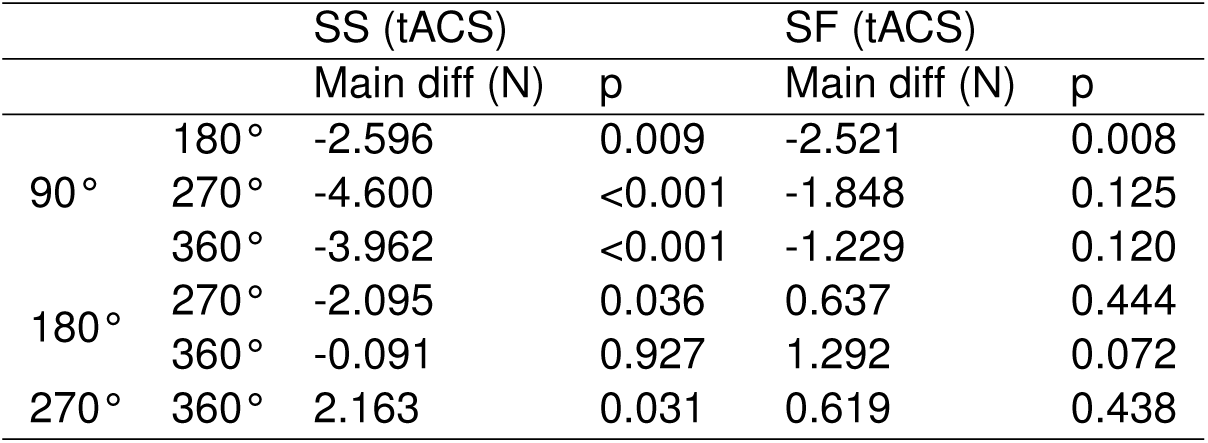
Post hoc analyses followed the Friedman test for SS and repeated-measures ANOVA for SF. No correction was applied. Bold values indicate statistically significant differences at *p <* 0.05. Significant differences were found in SS between 90*^◦^* and 180*^◦^*–270*^◦^*–360*^◦^*, 180*^◦^* and 270*^◦^*, and between 270*^◦^* and 360*^◦^* phases (Wilcoxon signed-rank tests). In SF, a significant difference was observed between 90*^◦^* and 180*^◦^*.

**Table S2.**
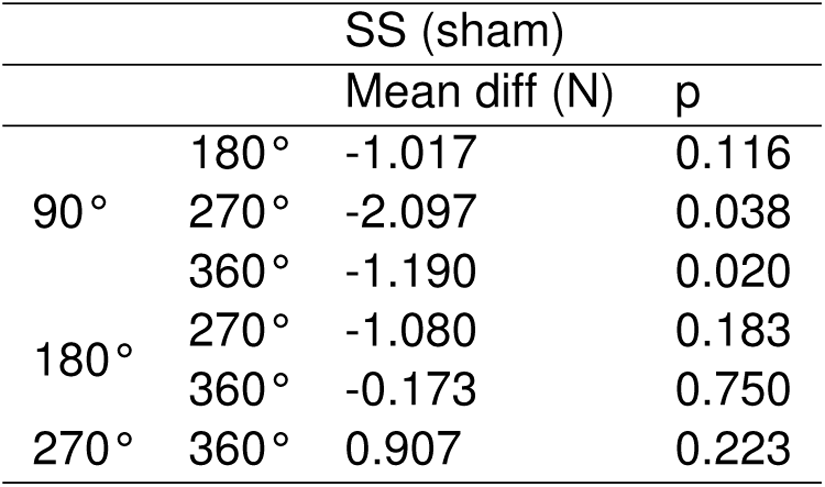
Post hoc analyses conducted after repeated-measures ANOVA. No correction was applied. Bold values indicate statistically significant differences at *p <* 0.05. Significant differences were found only in SS between 90*^◦^* and 270*^◦^*–360*^◦^*.

### Supplementary 2 – Phase Precession Pattern Using Multiple Window Sizes

Phase precession plots using different window sizes for experiment 1 (tACS) and 2 (sham) are shown in Figures S1 and S2. In the SS trials, the phase range was relatively small (60*^◦^*-120*^◦^*). The significant shift seen with larger time windows in this case is likely due to noise being smoothed out, rather than a meaningful underlying signal. On the other hand, in the SF trials, the phase shift was much larger (130*^◦^*-300*^◦^*), indicating a stronger effect than noise. Using large windows may have averaged out important circular changes, making the precession harder to detect. This interpretation is supported by a permutation test, which validates that the significance shift was only observed in Experiment 1-SF (Figure S3).

**Figure S1.**
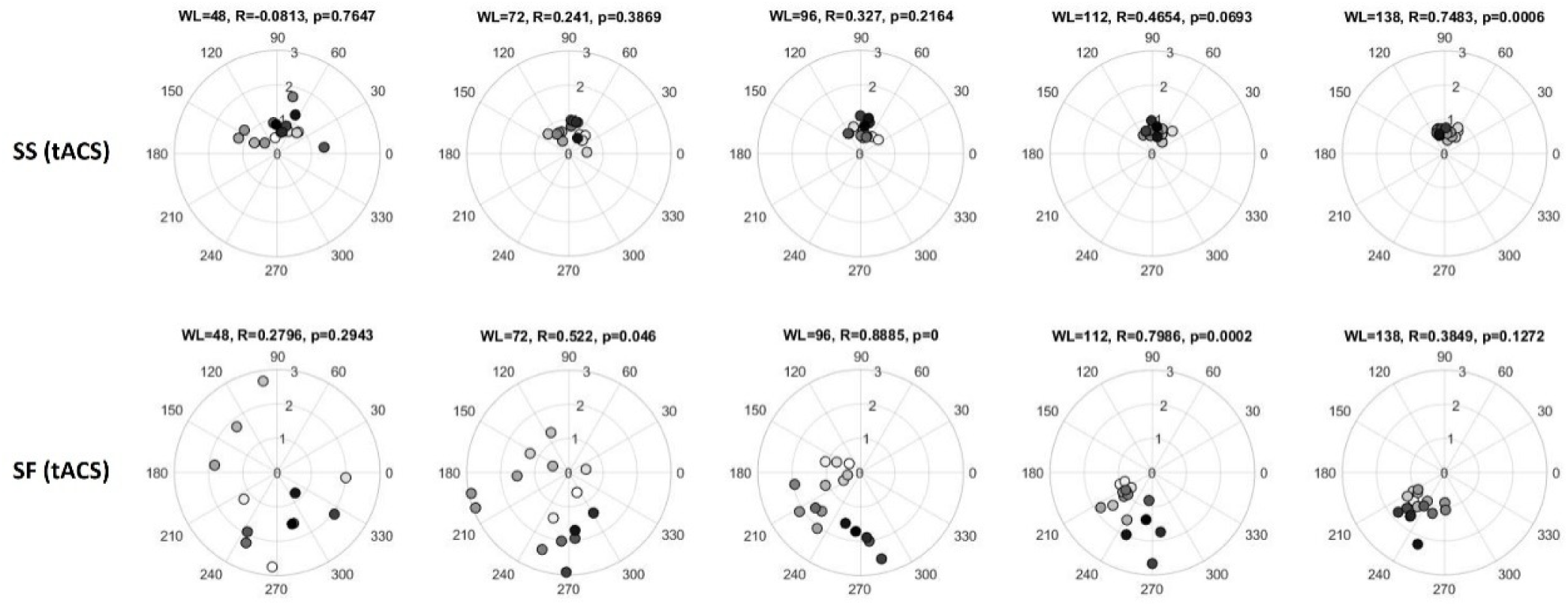
Experiment 1. Phase precession plots using different sliding window lengths (WL) with a fixed 16-step resolution. In some cases (e.g., WL = 48), fewer than 16 steps were available when, within a given window, one or more participants lacked SS or SF trials in every phase bin, making the phase estimation impossible.

**Figure S2.**
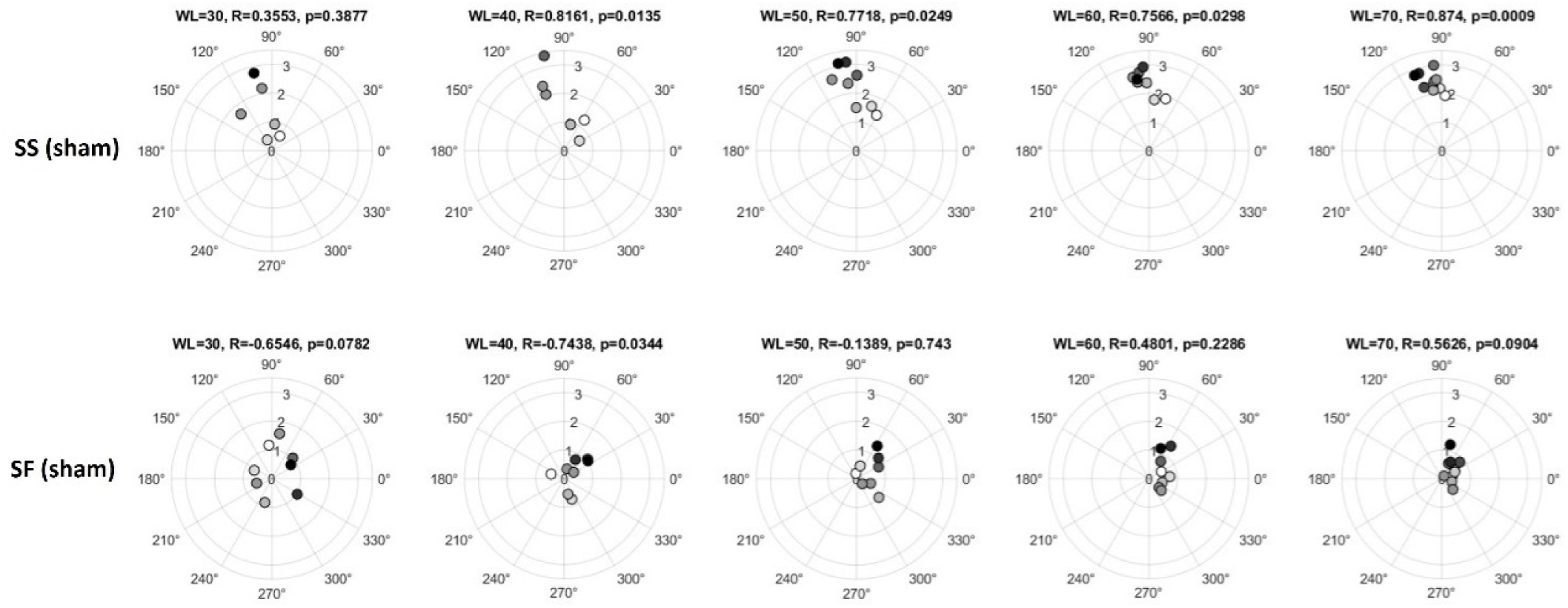
Experiment 2. Preferred phase precession with different sliding window lengths (WL) and a fixed 8-step resolution. In some cases, fewer than 8 steps were possible when, within a given window, one or more participants lacked SS or SF trials in any of the phase bins, making the phase estimation impossible.

### Supplementary 3 – Supplementary 3 – Permutation Test Results

**Figure S3.**
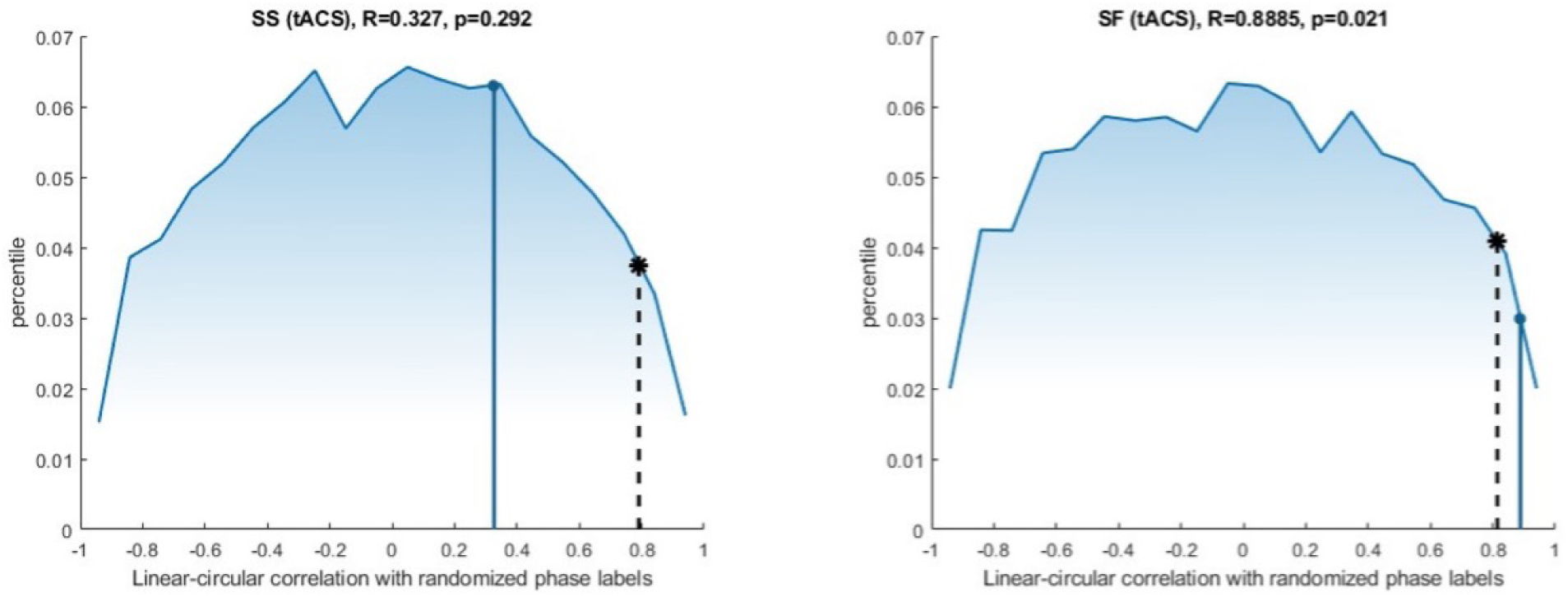
Permutation test with 10000 permutations randomizing the phase labels on linear-circular correlations between phases and force in the tACS group. The observed correlations are marked by the blue dots (*p* = 0.292 in SS (tACS) (left); *p* = 0.021 in SF (tACS) (right)). The black dashed lines with asterisks indicate the 95th percentile of the distribution. In SF (tACS) trials, the observed correlation laying beyond the 95th percentile, indicating a significant correlation between preferred phase and time points. In other trials, the observed correlation falls within the distribution, indicating a non-significant.

**Figure S4.**
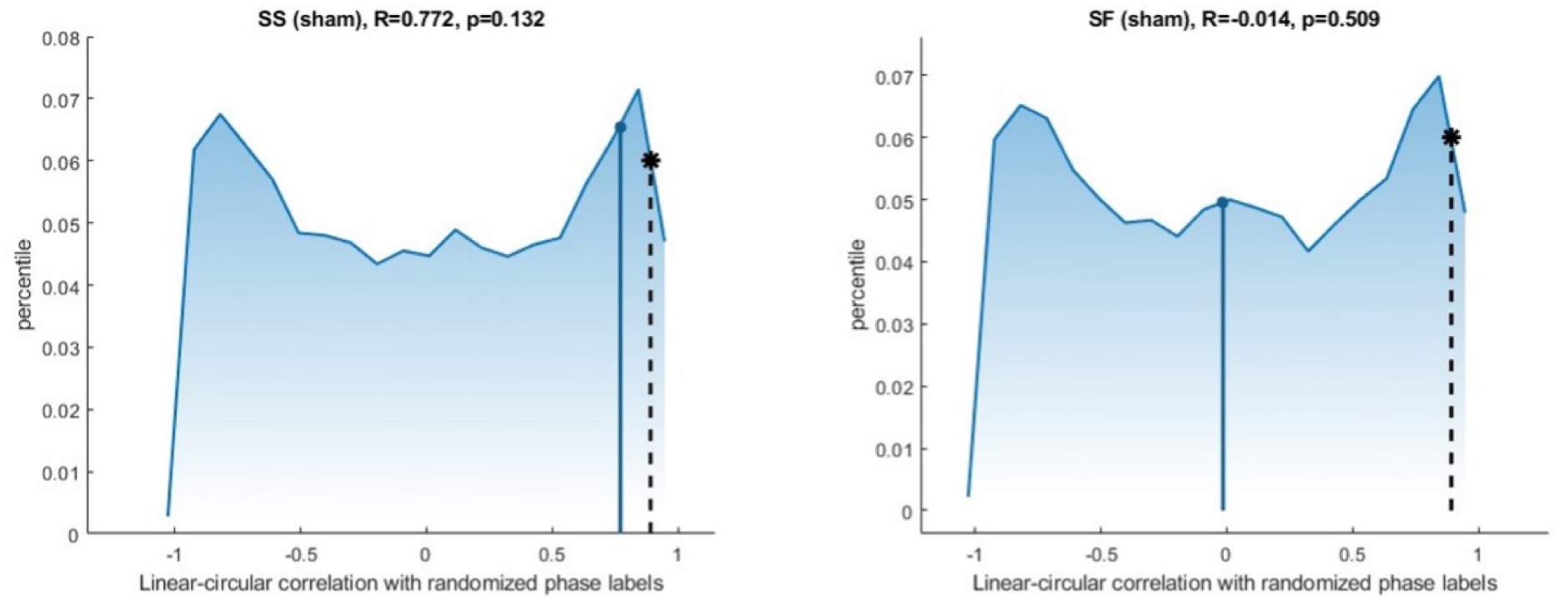
Permutation test with 10000 permutations randomizing the phase labels on linear-circular correlations between phases and force in the sham group. The observed correlations are marked by the blue dots (*p* = 0.132 in SS (sham) (left); *p* = 0.509 in SF (sham) (right)). The black dashed lines with asterisks indicate the 95th percentile of the distribution. Neither SS and SF of sham has passed the test.

