## Supplementary material for "tACS induces state-dependent shifts in inhibitory phase preference consistent with phase precession": Latex project: 00_Article_Merge.pdf

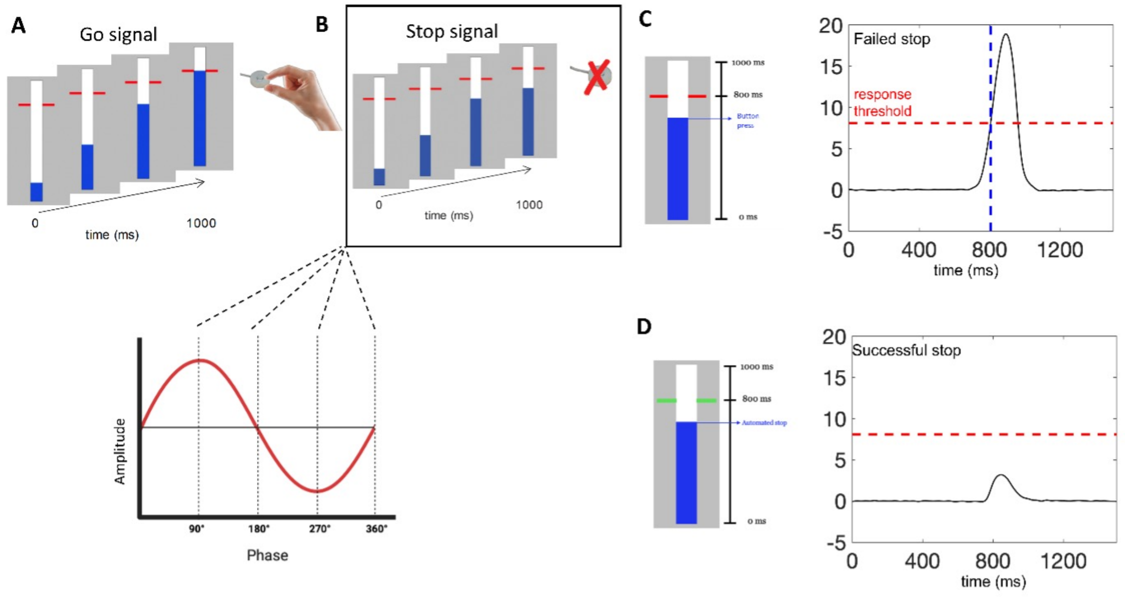


**Figure 1. Anticipatory stop-signal task and trial classification.**
(**A**) In the go trials, participants squeezed a force sensor as a bar reached the target. (**B**) In stop trials (30% of the total trials), the bar stopped early, requiring inhibition. In Experiment 1, the stop signals were presented equiprobably at four phase angles (90○, 180○, 270○, 360○) relative to the tACS; in Experiment 2, the instantaneous phase at the stop signals was estimated post hoc and assigned to the same four phase bins. (**C**) Stop-failed (SF) were defined as responses exceeding the response threshold. The target line turned red to indicate failed inhibition. (**D**) Stop-successful (SS) were those in which force remained below the threshold; the target line turned green to indicate successful inhibition.

phase at the instant corresponding to the original endpoint. Estimated phases were categorized into four bins: 90° ($\geq 45°$ to $< 135°$), 180° ($\geq 135°$ to $< 225°$), 270° ($\geq 225°$ to $< 315°$), and 360° ($\geq 315°$ to $< 360°$ or $\geq 0°$ to $< 45°$).

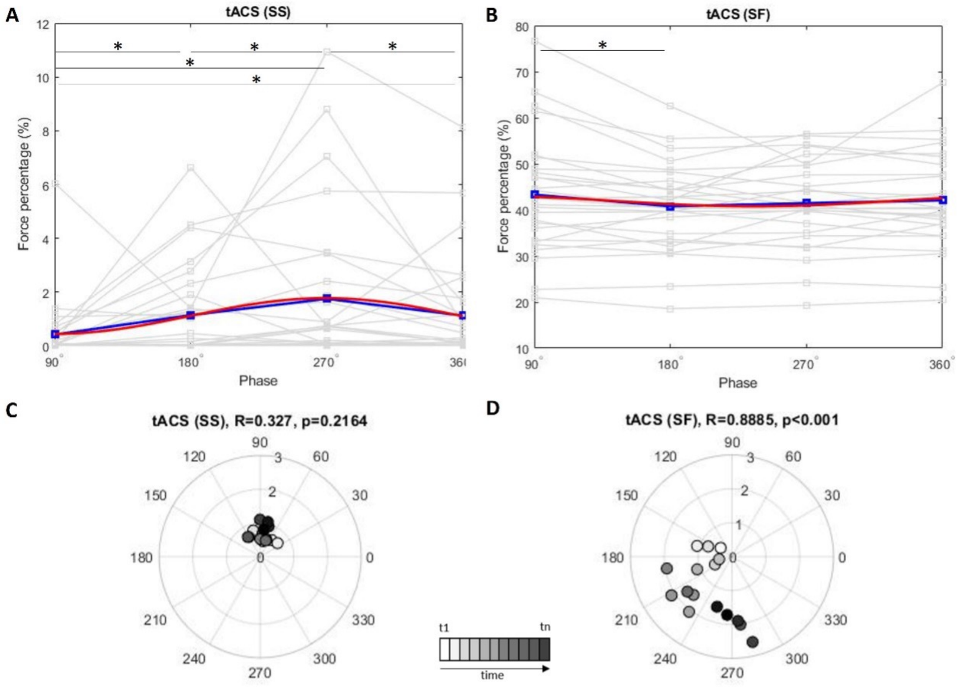


**Figure 2. Experiment 1**. (**A**)–(**B**), plots of peak force normalized with respect to the maximal voluntary force across tACS beta phases for stop success (SS) and stop failure (SF) trials. Gray lines represent individual data points. The blue lines indicate the average data, while the red curve shows the best-fitting sinusoidal pattern for each condition. Less force refers to greater inhibition. (**C**)–(**D**), polar plots of phase shifting of inhibition over time. Angular position represents preferred phase of inhibition, and radius represents the weight. A significant shift is observed in the SF group (~120° to ~300°, $R = 0.889$, $p < 0.001$), but not the SS ($R = 0.327$, $p = 0.216$).

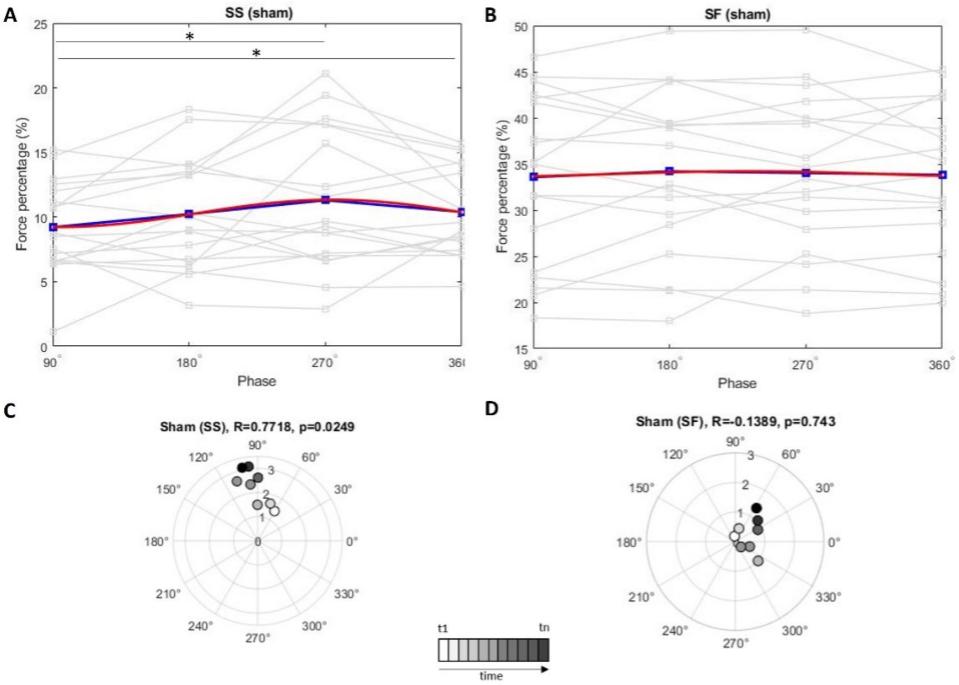


**Figure 3. Experiment 2**. (**A**)–(**B**), plots of normalized peak force across Fz-EEG beta phases for stop success (SS) and stop failure (SF) trials. Gray lines represent individual data points. The blue lines indicate the average data, while the red curve shows the best-fitting sinusoidal pattern for each condition. (**C**)–(**D**), polar plots of phase shifting of inhibition over time.

### Supplementary Information

#### Supplementary 1 - Post hoc Comparisons of Force Modulation by Beta Phase

| | | SS (tACS) | | SF (tACS) | |
|---|---|---|---|---|---|
| | | Main diff (N) | p | Main diff (N) | p |
| 90° | 180° | -2.596 | 0.009 | -2.521 | 0.008 |
| | 270° | -4.600 | <0.001 | -1.848 | 0.125 |
| | 360° | -3.962 | <0.001 | -1.229 | 0.120 |
| 180° | 270° | -2.095 | 0.036 | 0.637 | 0.444 |
| | 360° | -0.091 | 0.927 | 1.292 | 0.072 |
| 270° | 360° | 2.163 | 0.031 | 0.619 | 0.438 |

**Table S1.** Post hoc analyses followed the Friedman test for SS and repeated-measures ANOVA for SF. No correction was applied. Bold values indicate statistically significant differences at $p < 0.05$. Significant differences were found in SS between 90° and 180°–270°–360°, 180° and 270°, and between 270° and 360° phases (Wilcoxon signed-rank tests). In SF, a significant difference was observed between 90° and 180°.

| | | SS (sham) | |
|---|---|---|---|
| | | Mean diff (N) | p |
| 90° | 180° | -1.017 | 0.116 |
| | 270° | -2.097 | 0.038 |
| | 360° | -1.190 | 0.020 |
| 180° | 270° | -1.080 | 0.183 |
| | 360° | -0.173 | 0.750 |
| 270° | 360° | 0.907 | 0.223 |

**Table S2.** Post hoc analyses conducted after repeated-measures ANOVA. No correction was applied. Bold values indicate statistically significant differences at $p < 0.05$. Significant differences were found only in SS between 90° and 270°–360°.

#### Supplementary 2 - Phase Precession Pattern Using Multiple Window Sizes

Phase precession plots using different window sizes for experiment 1 (tACS) and 2 (sham) are shown in Figures S1 and S2. In the SS trials, the phase range was relatively small ($60°$-$120°$). The significant shift seen with larger time windows in this case is likely due to noise being smoothed out, rather than a meaningful underlying signal. On the other hand, in the SF trials, the phase shift was much larger ($130°$-$300°$), indicating a stronger effect than noise. Using large windows may have averaged out important circular changes, making the precession harder to detect. This interpretation is supported by a permutation test, which validates that the significance shift was only observed in Experiment 1-SF (Figure S3).

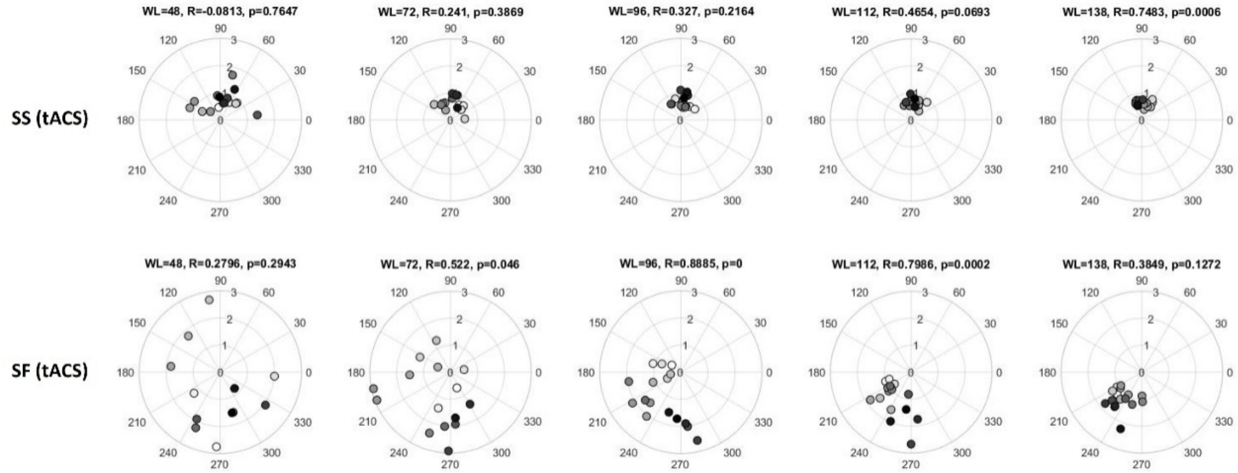


**Figure S1. Experiment 1.** Phase precession plots using different sliding window lengths (WL) with a fixed 16-step resolution. In some cases (e.g., WL = 48), fewer than 16 steps were available when, within a given window, one or more participants lacked SS or SF trials in every phase bin, making the phase estimation impossible.

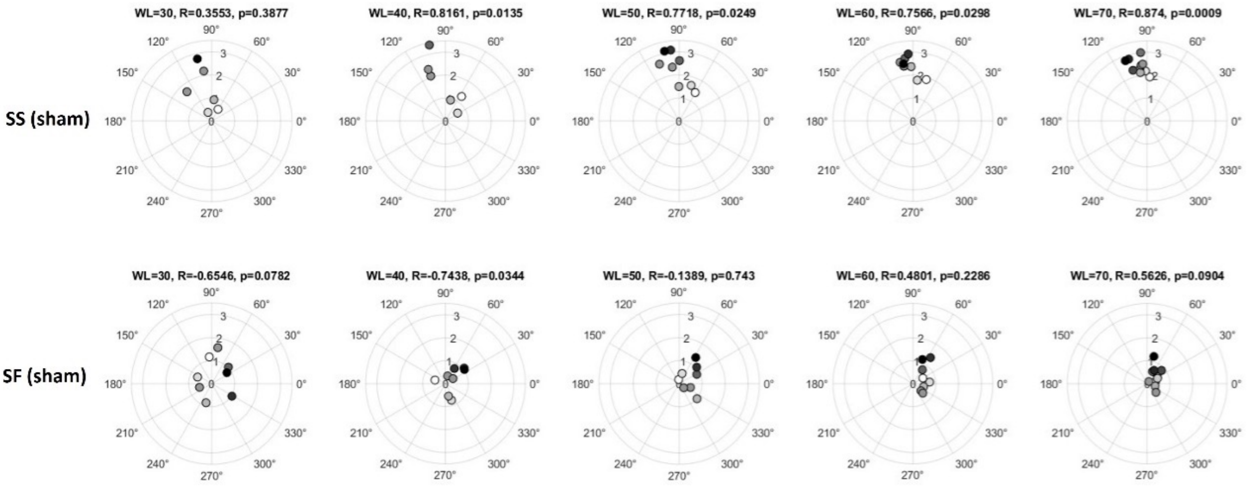


**Figure S2. Experiment 2.** Preferred phase precession with different sliding window lengths (WL) and a fixed 8-step resolution. In some cases, fewer than 8 steps were possible when, within a given window, one or more participants lacked SS or SF trials in any of the phase bins, making the phase estimation impossible.

#### Supplementary 3 - Supplementary 3 - Permutation Test Results

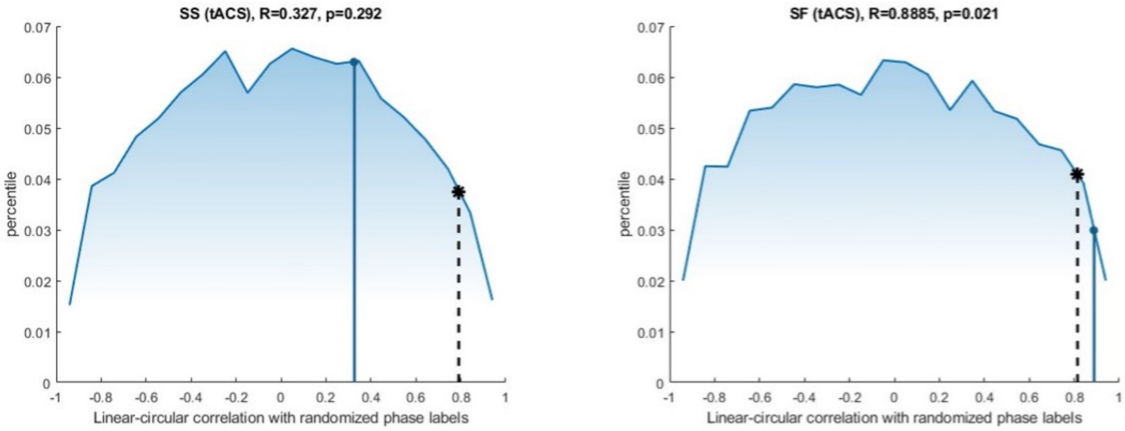


**Figure S3.** Permutation test with 10000 permutations randomizing the phase labels on linear-circular correlations between phases and force in the tACS group. The observed correlations are marked by the blue dots ($p = 0.292$ in SS (tACS) (left); $p = 0.021$ in SF (tACS) (right)). The black dashed lines with asterisks indicate the 95th percentile of the distribution. In SF (tACS) trials, the observed correlation laying beyond the 95th percentile, indicating a significant correlation between preferred phase and time points. In other trials, the observed correlation falls within the distribution, indicating a non-significant.

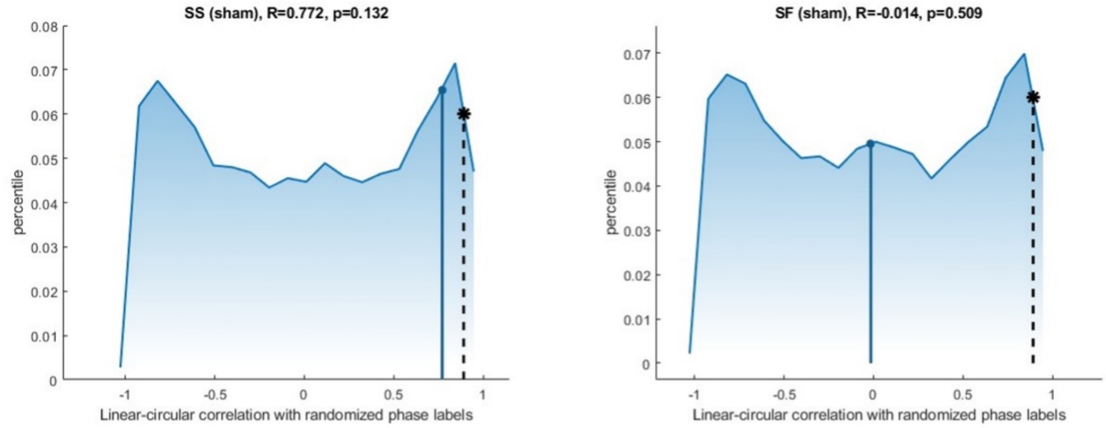


**Figure S4.** Permutation test with 10000 permutations randomizing the phase labels on linear-circular correlations between phases and force in the sham group. The observed correlations are marked by the blue dots ($p = 0.132$ in SS (sham) (left); $p = 0.509$ in SF (sham) (right)). The black dashed lines with asterisks indicate the 95th percentile of the distribution. Neither SS and SF of sham has passed the test.
