## Supplementary material for "tACS induces state-dependent shifts in inhibitory phase preference consistent with phase precession": Latex project: 00_Article_Merge.tex

%% select between bioRxiv style or submission style by commenting either line
%\documentclass[towcolumn]{bioRxiv}
\documentclass[submit]{bioRxiv}
\renewcommand\familydefault{\sfdefault}
\usepackage{lipsum}
\usepackage{wasysym}
\usepackage{hyperref}

%% setup specific to this preprint -- add or remove as you need.
\hyphenation{empire} % prevents word "empire" being split across lines
\sisetup{range-units = single,
        separate-uncertainty = true,
        multi-part-units = single,
        range-phrase = --} % units setup
\usepackage{multirow} % allows merging of cells in tables

\begin{document}

%% comment out any files that you don't want to include as sections
\include{01_Article_MainText}
\include{02_Article_Supplementary}

\end{document}
