## Supplementary material for "tACS induces state-dependent shifts in inhibitory phase preference consistent with phase precession": Latex project: 02_Article_Supplementary.tex

%  CONFIGURE NEW SINGLE-PAGE FORMAT 

\onecolumn % go back to one column
\fancyhead{} % make sure we get no headers
\renewcommand{\floatpagefraction}{0.1}
\lfoot[\bSupInf]{\dAuthor}
\rfoot[\dAuthor]{\cSupInf}
\newpage

\captionsetup*{format=largeformat} % make figure legend slightly larger than in the paper
\setcounter{figure}{0} % reset figure counter for Supp. Figures
\setcounter{equation}{0} % reset equation counter for Supp. Equations
%\setcounter{page}{1} % reset page count
\makeatletter 
\renewcommand{\thefigure}{S\@arabic\c@figure} % make Figure legend start with Figure S
\renewcommand{\thetable}{S\@arabic\c@table} % make Table legend start with Table S
\makeatother
\def\theequation{S\arabic{equation}}

%  MAIN TEXT 

\newpage
\section*{Supplementary Information}
\phantomsection
\label{supplementary}

\subsection*{Supplementary 1 - Post hoc Comparisons of Force Modulation by Beta Phase}

% Please add the following required packages to your document preamble:
% \usepackage{multirow}
\begin{table}[h]
    \title{\textbf{Post hoc comparisons of force modulation by beta phase in Experiment 1 (tACS)}}

\begin{tabular}{llllll}
\hline
                      &      & \multicolumn{2}{l}{SS (tACS)}    & \multicolumn{2}{l}{SF (tACS)} \\ \hline
                      &      & Main diff (N) & p                & Main diff (N)     & p         \\ \hline
\multirow{3}{*}{90°}  & 180° & -2.596        & 0.009            & -2.521            & 0.008     \\
                      & 270° & -4.600        & \textless{}0.001 & -1.848            & 0.125     \\
                      & 360° & -3.962        & \textless{}0.001 & -1.229            & 0.120     \\
\multirow{2}{*}{180°} & 270° & -2.095        & 0.036            & 0.637             & 0.444     \\
                      & 360° & -0.091        & 0.927            & 1.292             & 0.072     \\
270°                  & 360° & 2.163         & 0.031            & 0.619             & 0.438     \\ \hline
\end{tabular}
\caption{Post hoc analyses followed the Friedman test for SS and repeated-measures ANOVA for SF. No correction was applied. Bold values indicate statistically significant differences at $p<0.05$. Significant differences were found in SS between 90$^{\circ}$ and 180$^{\circ}$--270$^{\circ}$--360$^{\circ}$, 180$^{\circ}$ and 270$^{\circ}$, and between 270$^{\circ}$ and 360$^{\circ}$ phases (Wilcoxon signed-rank tests). In SF, a significant difference was observed between 90$^{\circ}$ and 180$^{\circ}$.
}
    \label{reftab:t1}
\end{table}

% Please add the following required packages to your document preamble:
% \usepackage{multirow}
\begin{table}[h]
    \title{\textbf{Post hoc comparisons of force modulation by beta phase in Experiment 2 (sham)}}
\begin{tabular}{llll}
\hline
                      &      & \multicolumn{2}{l}{SS (sham)} \\ \hline
                      &      & Mean diff (N)     & p         \\ \hline
\multirow{3}{*}{90°}  & 180° & -1.017            & 0.116     \\
                      & 270° & -2.097            & 0.038     \\
                      & 360° & -1.190            & 0.020     \\
\multirow{2}{*}{180°} & 270° & -1.080            & 0.183     \\
                      & 360° & -0.173            & 0.750     \\
270°                  & 360° & 0.907             & 0.223     \\ \hline
\end{tabular}
\caption{Post hoc analyses conducted after repeated-measures ANOVA. No correction was applied. Bold values indicate statistically significant differences at $p<0.05$. Significant differences were found only in SS between 90$^{\circ}$ and 270$^{\circ}$--360$^{\circ}$.
}
    \label{reftab:t2}
\end{table}

\subsection*{Supplementary 2 - Phase Precession Pattern Using Multiple Window Sizes}

Phase precession plots using different window sizes for experiment 1 (tACS) and 2 (sham) are shown in Figures S1 and S2. In the SS trials, the phase range was relatively small (\( 60^\circ\)-\( 120^\circ \)). The significant shift seen with larger time windows in this case is likely due to noise being smoothed out, rather than a meaningful underlying signal. On the other hand, in the SF trials, the phase shift was much larger (\( 130^\circ\)-\( 300^\circ \)), indicating a stronger effect than noise. Using large windows may have averaged out important circular changes, making the precession harder to detect. 

This interpretation is supported by a permutation test, which validates that the significance shift was only observed in Experiment 1-SF (Figure S3).

\begin{figure}[h]
    \centering
    \includegraphics[width=1.1\textwidth]{Figures/figs1.jpg}
    \caption{\textbf{Experiment 1.} Phase precession plots using different sliding window lengths (WL) with a fixed 16-step resolution. In some cases (e.g., WL = 48), fewer than 16 steps were available when, within a given window, one or more participants lacked SS or SF trials in every phase bin, making the phase estimation impossible.}
    \label{reffig:s1}
\end{figure}

\begin{figure}[h]
    \centering
    \includegraphics[width=1.1\textwidth]{Figures/figs2.jpg}
    \caption{\textbf{Experiment 2.} Preferred phase precession with different sliding window lengths (WL) and a fixed 8-step resolution. In some cases, fewer than 8 steps were possible when, within a given window, one or more participants lacked SS or SF trials in any of the phase bins, making the phase estimation impossible.}
    \label{reffig:s2}
\end{figure}

\subsection*{Supplementary 3 - Supplementary 3 - Permutation Test Results}

\begin{figure}[h]
    \centering
    \includegraphics[width=0.9\textwidth]{Figures/figs3.jpg}
    \caption{Permutation test with 10000 permutations randomizing the phase labels on linear-circular correlations between phases and force in the tACS group. The observed correlations are marked by the blue dots (\( p = 0.292 \) in SS (tACS) (left); \( p = 0.021 \) in SF (tACS) (right)). The black dashed lines with asterisks indicate the 95th percentile of the distribution. In SF (tACS) trials, the observed correlation laying beyond the 95th percentile, indicating a significant correlation between preferred phase and time points. In other trials, the observed correlation falls within the distribution, indicating a non-significant.}
    \label{reffig:s3}
\end{figure}    

\begin{figure}
    \centering
    \includegraphics[width=0.9\textwidth]{Figures/figs4.jpg}
    \caption{Permutation test with 10000 permutations randomizing the phase labels on linear-circular correlations between phases and force in the sham group. The observed correlations are marked by the blue dots (\( p = 0.132 \) in SS (sham) (left); \( p = 0.509 \) in SF (sham) (right)). The black dashed lines with asterisks indicate the 95th percentile of the distribution. Neither SS and SF of sham has passed the test.}
    \label{reffig:s4}
\end{figure}
