## Supplementary material for "tACS induces state-dependent shifts in inhibitory phase preference consistent with phase precession": Latex project: README.md

# manuscript-templates
LaTeX templates for manuscripts (switchable bioRxiv preprint and journal submission templates), for use on Overleaf.

## Basic idea

An ideal manuscript LaTeX template should be able to generate two versions of the manuscript:

1. a preprint version - typeset to look like a finished paper
2. a journal submission version - better for reviewing, with line numbers and different spacing, etc.

The ideal template should be able to switch between these two manuscript layouts. A preview of both outputs is shown below.

## Use

To use, just clone this repo and import into Overleaf or a directory on your own computer to use a local LaTeX installation.
The following files and folders are not required: `img`, `Examples`, `README`.

Write your manuscript in `01_Article_MainText.tex` and then comment either line 2 or 3 of `00_Article_Merge.tex` to select between outputs.
There are optional Supplementary tex files that can also be edited.
If they are not required, comment the appropriate lines in `00_Article_Merge.tex`.

Tested on Overleaf - TeXLive 2023 with pdfLaTeX compiler.


## Contributions

The initial overleaf template was forked from the template created by Henriques lab [available here](https://www.overleaf.com/latex/templates/henriqueslab-biorxiv-template/nyprsybwffws).
This is a modification of the PNAS template (also available on Overleaf) which is widely used on bioRxiv.
We have made a few changes to the HenriquesLab bioRxiv template and made it possible to easily generate a journal submission version.

Changes include:

- genuine bioRxiv logo used in the footer
- support for ORCiDs using `\orcidlink`
- Helvetica for readability
- greater range of custom units
- refactored into one single class - thank you to Dominik Straub for his work on [this fork](https://github.com/dominikstrb/manuscript-templates).

## What do they look like?

### Preprint version (for bioRxiv)

![img](img/Example_bioRxiv.png?raw=true "image")

### Journal submission version

![img](img/Example_submit.png?raw=true "image")
