## Supplementary material for "tACS induces state-dependent shifts in inhibitory phase preference consistent with phase precession": Latex project: refs.bib

@article{Fang2024,
  author  = {Fang, Zhou and ten Oever, Sanne and Sack, Alexander T. and Leunissen, Inge},
  title   = {tACS induces state-dependent shifts in inhibitory phase preference consistent with phase precession},
  journal = {NeuroImage},
  year    = {2024},
  note    = {Advance online / in press; preprint on bioRxiv: 10.1101/2023.09.19.558205},
}

@article{Krause2019,
  title = {Transcranial alternating current stimulation entrains single-neuron activity in the primate brain},
  volume = {116},
  ISSN = {1091-6490},
  url = {http://dx.doi.org/10.1073/pnas.1815958116},
  DOI = {10.1073/pnas.1815958116},
  number = {12},
  journal = {Proceedings of the National Academy of Sciences},
  publisher = {Proceedings of the National Academy of Sciences},
  author = {Krause,  Matthew R. and Vieira,  Pedro G. and Csorba,  Bennett A. and Pilly,  Praveen K. and Pack,  Christopher C.},
  year = {2019},
  month = mar,
  pages = {5747–5755}
}

@article{Buzsaki2004,
  author  = {Buzsaki, Gyorgy and Draguhn, Andreas},
  title   = {Neuronal oscillations in cortical networks},
  journal = {Science},
  year    = {2004},
  volume  = {304},
  number  = {5679},
  pages   = {1926--1929},
  doi     = {10.1126/science.1099745}
}

@article{OKeefe1993,
  author  = {O'Keefe, John and Recce, Michael L.},
  title   = {Phase relationship between hippocampal place units and the {EEG} theta rhythm},
  journal = {Hippocampus},
  year    = {1993},
  volume  = {3},
  number  = {3},
  pages   = {317--330},
  doi     = {10.1002/hipo.450030307}
}

@article{vanderMeer2011,
  author  = {van der Meer, Matthijs A. A. and Redish, A. David},
  title   = {Theta phase precession in rat ventral striatum links place and reward information},
  journal = {Journal of Neuroscience},
  year    = {2011},
  volume  = {31},
  number  = {8},
  pages   = {2843--2854},
  doi     = {10.1523/JNEUROSCI.4869-10.2011}
}

@article{Jones2005,
  author  = {Jones, Matthew W. and Wilson, Matthew A.},
  title   = {Phase precession of medial prefrontal cortical activity relative to the hippocampal theta rhythm},
  journal = {Hippocampus},
  year    = {2005},
  volume  = {15},
  number  = {7},
  pages   = {867--873},
  doi     = {10.1002/hipo.20119}
}

@article{Lisman2013,
  author  = {Lisman, John E. and Jensen, Ole},
  title   = {The theta--gamma neural code},
  journal = {Neuron},
  year    = {2013},
  volume  = {77},
  number  = {6},
  pages   = {1002--1016},
  doi     = {10.1016/j.neuron.2013.03.007}
}

@article{Widloski2014,
  author  = {Widloski, John and Fiete, Ila R.},
  title   = {A model of grid cell development through spatial exploration and spike time-dependent plasticity},
  journal = {Neuron},
  year    = {2014},
  volume  = {83},
  number  = {2},
  pages   = {481--495},
  doi     = {10.1016/j.neuron.2014.06.018}
}

@article{Buzsaki2018,
  author  = {Buzsaki, Gyorgy and Tingley, David},
  title   = {Space and time: The hippocampus as a sequence generator},
  journal = {Trends in Cognitive Sciences},
  year    = {2018},
  volume  = {22},
  number  = {10},
  pages   = {853--869},
  doi     = {10.1016/j.tics.2018.07.006}
}

@article{Qasim2021,
  author  = {Qasim, Salman E. and Fried, Itzhak and Jacobs, Joshua},
  title   = {Phase precession in the human hippocampus and entorhinal cortex},
  journal = {Cell},
  year    = {2021},
  volume  = {184},
  number  = {12},
  pages   = {3242--3255.e10},
  doi     = {10.1016/j.cell.2021.04.017}
}

@article{Reddy2021,
  author  = {Reddy, Leila and Self, Matthew W. and Zoefel, Benedikt and Poncet, Marlene and Possel, Jessy K. and Peters, Judith C. and Baayen, Johannes C. and Idema, Sander and VanRullen, Rufin and Roelfsema, Pieter R.},
  title   = {Theta-phase dependent neuronal coding during sequence learning in human single neurons},
  journal = {Nature Communications},
  year    = {2021},
  volume  = {12},
  number  = {1},
  pages   = {4839},
  doi     = {10.1038/s41467-021-25150-0}
}

@article{Ozen2010,
  author  = {Ozen, Simal and Sirota, Anton and Belluscio, Mariano A. and Anastassiou, Costas A. and Stark, Eran and Koch, Christof and Buzsaki, Gyorgy},
  title   = {Transcranial electric stimulation entrains cortical neuronal populations in rats},
  journal = {Journal of Neuroscience},
  year    = {2010},
  volume  = {30},
  number  = {34},
  pages   = {11476--11485},
  doi     = {10.1523/JNEUROSCI.5252-09.2010}
}

@article{Johnson2020,
  author  = {Johnson, Leah and Alekseichuk, Ivan and Krieg, Johannes and Doyle, Andrew and Yu, Yan and Vitek, Jan and Özen, Simal and Frohlich, Flavio},
  title   = {Dose-dependent effects of transcranial alternating current stimulation on spike timing in awake nonhuman primates},
  journal = {Science Advances},
  year    = {2020},
  volume  = {6},
  number  = {36},
  pages   = {eaaz2747},
  doi     = {10.1126/sciadv.aaz2747}
}

@article{Wischnewski2019,
  author  = {Wischnewski, Miles and Schutter, Dennis J. L. G. and Nitsche, Michael A.},
  title   = {Effects of beta-tACS on corticospinal excitability: A meta-analysis},
  journal = {Brain Stimulation},
  year    = {2019},
  volume  = {12},
  number  = {6},
  pages   = {1381--1389},
  doi     = {10.1016/j.brs.2019.07.023}
}

@article{Schilberg2018,
  author  = {Schilberg, Lukas and Engelen, Tahnee and ten Oever, Sanne and Schuhmann, Teresa and de Gelder, Beatrice and de Graaf, Tom A. and Sack, Alexander T.},
  title   = {Phase of beta-frequency tACS over primary motor cortex modulates corticospinal excitability},
  journal = {Cortex},
  year    = {2018},
  volume  = {103},
  pages   = {142--152},
  doi     = {10.1016/j.cortex.2018.03.001}
}

@article{Bikson2009,
  author  = {Bikson, Marom and Datta, Abhishek and Elwassif, Maged},
  title   = {Establishing safety limits for transcranial direct current stimulation},
  journal = {Clinical Neurophysiology},
  year    = {2009},
  volume  = {120},
  number  = {6},
  pages   = {1033--1034},
  doi     = {10.1016/j.clinph.2009.03.018}
}

@article{Woods2016,
  author  = {Woods, Alexander J. and Antal, Andrea and Bikson, Marom and Boggio, Paulo S. and Brunoni, Andre R. and Celnik, Pablo and Cohen, Leonardo G. and Fregni, Felipe and Herrmann, Christoph S. and Kappenman, Emily S. and Knotkova, Hana and Liebetanz, David and Miniussi, Carlo and Miranda, Pedro C. and Paulus, Walter and Priori, Alberto and Reato, Davide and Stagg, Charlotte and Wenderoth, Nicole and Nitsche, Michael A.},
  title   = {A technical guide to tDCS, and related non-invasive brain stimulation tools},
  journal = {Clinical Neurophysiology},
  year    = {2016},
  volume  = {127},
  number  = {2},
  pages   = {1031--1048},
  doi     = {10.1016/j.clinph.2015.11.012}
}

@article{Oldfield1971,
  author  = {Oldfield, R. C.},
  title   = {The assessment and analysis of handedness: The {Edinburgh} inventory},
  journal = {Neuropsychologia},
  year    = {1971},
  volume  = {9},
  number  = {1},
  pages   = {97--113},
  doi     = {10.1016/0028-3932(71)90067-4}
}

@article{Pogosyan2009,
  author  = {Pogosyan, Alex and Gaynor, Louise D. and Eusebio, Alexandre and Brown, Peter},
  title   = {Boosting cortical activity at beta-band frequencies slows movement in humans},
  journal = {Current Biology},
  year    = {2009},
  volume  = {19},
  number  = {19},
  pages   = {1637--1641},
  doi     = {10.1016/j.cub.2009.07.074}
}

@article{Dan2006,
  author  = {Dan, Yang and Poo, Mu-ming},
  title   = {Spike timing-dependent plasticity: From synapse to perception},
  journal = {Physiological Reviews},
  year    = {2006},
  volume  = {86},
  number  = {3},
  pages   = {1033--1048},
  doi     = {10.1152/physrev.00030.2005}
}

@article{Lengyel2005,
  author  = {Lengyel, Mate and Kwag, Jeehyun and Paulsen, Ole and Dayan, Peter},
  title   = {Matching storage and recall: Hippocampal spike timing-dependent plasticity and phase response curves},
  journal = {Nature Neuroscience},
  year    = {2005},
  volume  = {8},
  number  = {12},
  pages   = {1677--1683},
  doi     = {10.1038/nn1561}
}

@article{Samsonovich1997,
  author  = {Samsonovich, Alexei and McNaughton, Bruce L.},
  title   = {Path integration and cognitive mapping in a continuous attractor neural network model},
  journal = {Journal of Neuroscience},
  year    = {1997},
  volume  = {17},
  number  = {15},
  pages   = {5900--5920},
  doi     = {10.1523/JNEUROSCI.17-15-05900.1997}
}

@article{Mehta2002,
  author  = {Mehta, Mayank R. and Lee, Albert K. and Wilson, Matthew A.},
  title   = {Role of experience and oscillations in transforming a rate code into a temporal code},
  journal = {Nature},
  year    = {2002},
  volume  = {417},
  number  = {6890},
  pages   = {741--746},
  doi     = {10.1038/nature00807}
}

@book{Kuramoto1984,
  author    = {Kuramoto, Yoshiki},
  title     = {Chemical Oscillations, Waves, and Turbulence},
  publisher = {Springer},
  address   = {Berlin},
  year      = {1984},
  doi       = {10.1007/978-3-642-69689-3}
}

@article{Oostenveld2011,
  author  = {Oostenveld, Robert and Fries, Pascal and Maris, Eric and Schoffelen, Jan-Mathijs},
  title   = {FieldTrip: Open source software for advanced analysis of {MEG}, {EEG}, and invasive electrophysiological data},
  journal = {Computational Intelligence and Neuroscience},
  year    = {2011},
  volume  = {2011},
  pages   = {156869},
  doi     = {10.1155/2011/156869}
}

@article{tenOever2020,
  author  = {ten Oever, Sanne and de Weerd, Peter and Sack, Alexander T.},
  title   = {Phase-dependent amplification of working memory content and performance},
  journal = {Nature Communications},
  year    = {2020},
  volume  = {11},
  number  = {1},
  pages   = {1832},
  doi     = {10.1038/s41467-020-15629-7}
}
@article{Jensen2005,
  author  = {Jensen, Ole and Lisman, John E.},
  title   = {Hippocampal sequence-encoding driven by a cortical multi-item working memory buffer},
  journal = {Trends in Neurosciences},
  year    = {2005},
  volume  = {28},
  number  = {2},
  pages   = {67--72},
  doi     = {10.1016/j.tins.2004.11.006}
}
@article{Helfrich2014,
  author  = {Helfrich, Randolph F. and Schneider, Thomas R. and Rach, Stefan and Trautmann-Lengsfeld, Sina A. and Engel, Andreas K. and Herrmann, Christoph S.},
  title   = {Entrainment of brain oscillations by transcranial alternating current stimulation},
  journal = {Current Biology},
  year    = {2014},
  volume  = {24},
  number  = {3},
  pages   = {333--339},
  doi     = {10.1016/j.cub.2013.12.041}
}

@article{Beste2023,
  title = {Towards a systematization of brain oscillatory activity in actions},
  volume = {6},
  ISSN = {2399-3642},
  url = {http://dx.doi.org/10.1038/s42003-023-04531-9},
  DOI = {10.1038/s42003-023-04531-9},
  number = {1},
  journal = {Communications Biology},
  publisher = {Springer Science and Business Media LLC},
  author = {Beste,  Christian and M\"{u}nchau,  Alexander and Frings,  Christian},
  year = {2023},
  month = feb 
}

@article{Thut2011,
  title = {Entrainment of Perceptually Relevant Brain Oscillations by Non-Invasive Rhythmic Stimulation of the Human Brain},
  volume = {2},
  ISSN = {1664-1078},
  url = {http://dx.doi.org/10.3389/fpsyg.2011.00170},
  DOI = {10.3389/fpsyg.2011.00170},
  journal = {Frontiers in Psychology},
  publisher = {Frontiers Media SA},
  author = {Thut,  Gregor and Schyns,  Philippe G. and Gross,  Joachim},
  year = {2011}
}

@article{Herrmann2016,
  title = {Shaping Intrinsic Neural Oscillations with Periodic Stimulation},
  volume = {36},
  ISSN = {1529-2401},
  url = {http://dx.doi.org/10.1523/JNEUROSCI.0236-16.2016},
  DOI = {10.1523/jneurosci.0236-16.2016},
  number = {19},
  journal = {The Journal of Neuroscience},
  publisher = {Society for Neuroscience},
  author = {Herrmann,  Christoph S. and Murray,  Micah M. and Ionta,  Silvio and Hutt,  Axel and Lefebvre,  Jérémie},
  year = {2016},
  month = may,
  pages = {5328–5337}
}

@article{Magee2001,
  title = {Dendritic Mechanisms of Phase Precession in Hippocampal CA1 Pyramidal Neurons},
  volume = {86},
  ISSN = {1522-1598},
  url = {http://dx.doi.org/10.1152/jn.2001.86.1.528},
  DOI = {10.1152/jn.2001.86.1.528},
  number = {1},
  journal = {Journal of Neurophysiology},
  publisher = {American Physiological Society},
  author = {Magee,  Jeffrey C.},
  year = {2001},
  month = jul,
  pages = {528–532}
}

@article{Narayanan2008,
  title = {The h Channel Mediates Location Dependence and Plasticity of Intrinsic Phase Response in Rat Hippocampal Neurons},
  volume = {28},
  ISSN = {1529-2401},
  url = {http://dx.doi.org/10.1523/JNEUROSCI.0835-08.2008},
  DOI = {10.1523/jneurosci.0835-08.2008},
  number = {22},
  journal = {The Journal of Neuroscience},
  publisher = {Society for Neuroscience},
  author = {Narayanan,  Rishikesh and Johnston,  Daniel},
  year = {2008},
  month = may,
  pages = {5846–5860}
}

@article{Wischnewski2024,
  title = {Induced neural phase precession through exogenous electric fields},
  volume = {15},
  ISSN = {2041-1723},
  url = {http://dx.doi.org/10.1038/s41467-024-45898-5},
  DOI = {10.1038/s41467-024-45898-5},
  number = {1},
  journal = {Nature Communications},
  publisher = {Springer Science and Business Media LLC},
  author = {Wischnewski,  Miles and Tran,  Harry and Zhao,  Zhihe and Shirinpour,  Sina and Haigh,  Zachary J. and Rotteveel,  Jonna and Perera,  Nipun D. and Alekseichuk,  Ivan and Zimmermann,  Jan and Opitz,  Alexander},
  year = {2024},
  month = feb 
}
@article{Schlesiger2015,
  title = {The medial entorhinal cortex is necessary for temporal organization of hippocampal neuronal activity},
  volume = {18},
  ISSN = {1546-1726},
  url = {http://dx.doi.org/10.1038/nn.4056},
  DOI = {10.1038/nn.4056},
  number = {8},
  journal = {Nature Neuroscience},
  publisher = {Springer Science and Business Media LLC},
  author = {Schlesiger,  Magdalene I and Cannova,  Christopher C and Boublil,  Brittney L and Hales,  Jena B and Mankin,  Emily A and Brandon,  Mark P and Leutgeb,  Jill K and Leibold,  Christian and Leutgeb,  Stefan},
  year = {2015},
  month = jun,
  pages = {1123–1132}
}

@article{SlaterHammel1960,
  title = {Reliability,  Accuracy,  and Refractoriness of a Transit Reaction},
  volume = {31},
  ISSN = {1067-1188},
  url = {http://dx.doi.org/10.1080/10671188.1960.10613098},
  DOI = {10.1080/10671188.1960.10613098},
  number = {2},
  journal = {Research Quarterly. American Association for Health,  Physical Education and Recreation},
  publisher = {Informa UK Limited},
  author = {Slater-Hammel,  A. T.},
  year = {1960},
  month = may,
  pages = {217–228}
}

@article{Coxon2006,
  title = {Intracortical Inhibition During Volitional Inhibition of Prepared Action},
  volume = {95},
  ISSN = {1522-1598},
  url = {http://dx.doi.org/10.1152/jn.01334.2005},
  DOI = {10.1152/jn.01334.2005},
  number = {6},
  journal = {Journal of Neurophysiology},
  publisher = {American Physiological Society},
  author = {Coxon,  James P. and Stinear,  Cathy M. and Byblow,  Winston D.},
  year = {2006},
  month = jun,
  pages = {3371–3383}
}

@article{Leunissen2017,
  title = {Reliable estimation of inhibitory efficiency: to anticipate,  choose or simply react?},
  volume = {45},
  ISSN = {1460-9568},
  url = {http://dx.doi.org/10.1111/ejn.13590},
  DOI = {10.1111/ejn.13590},
  number = {12},
  journal = {European Journal of Neuroscience},
  publisher = {Wiley},
  author = {Leunissen,  Inge and Zandbelt,  Bram B. and Potocanac,  Zrinka and Swinnen,  Stephan P. and Coxon,  James P.},
  editor = {Rossell,  Susan},
  year = {2017},
  month = jun,
  pages = {1512–1523}
}

@article{tenOever2016,
  title = {Stimulus Presentation at Specific Neuronal Oscillatory Phases Experimentally Controlled with tACS: Implementation and Applications},
  volume = {10},
  ISSN = {1662-5102},
  url = {http://dx.doi.org/10.3389/fncel.2016.00240},
  DOI = {10.3389/fncel.2016.00240},
  journal = {Frontiers in Cellular Neuroscience},
  publisher = {Frontiers Media SA},
  author = {ten Oever,  Sanne and de Graaf,  Tom A. and Bonnemayer,  Charlie and Ronner,  Jacco and Sack,  Alexander T. and Riecke,  Lars},
  year = {2016},
  month = oct 
}

@article{Cai2014,
author = {Cai, Weidong and Cannistraci, Christopher J. and Gore, John C. and Leung, Hoi-Chung},
title = {Sensorimotor-independent prefrontal activity during response inhibition},
journal = {Human Brain Mapping},
volume = {35},
number = {5},
pages = {2119-2136},
keywords = {prefrontal cortex, human, cognitive control, stop-signal task, fMRI},
doi = {https://doi.org/10.1002/hbm.22315},
url = {https://onlinelibrary.wiley.com/doi/abs/10.1002/hbm.22315},
year = {2014}
}

@article{VanHoornweder2025,
  title = {The causal role of beta band desynchronization: Individualized high-definition transcranial alternating current stimulation improves bimanual motor control},
  volume = {312},
  ISSN = {1053-8119},
  url = {http://dx.doi.org/10.1016/j.neuroimage.2025.121222},
  DOI = {10.1016/j.neuroimage.2025.121222},
  journal = {NeuroImage},
  publisher = {Elsevier BV},
  author = {Van Hoornweder,  Sybren and Mora,  Diego Andres Blanco and Nuyts,  Marten and Cuypers,  Koen and Verstraelen,  Stefanie and Meesen,  Raf},
  year = {2025},
  month = may,
  pages = {121222}
}

@article{Leunissen2022,
  title = {Effects of beta-band and gamma-band rhythmic stimulation on motor inhibition},
  volume = {25},
  ISSN = {2589-0042},
  url = {http://dx.doi.org/10.1016/j.isci.2022.104338},
  DOI = {10.1016/j.isci.2022.104338},
  number = {5},
  journal = {iScience},
  publisher = {Elsevier BV},
  author = {Leunissen,  Inge and Van Steenkiste,  Manon and Heise,  Kirstin-Friederike and Monteiro,  Thiago Santos and Dunovan,  Kyle and Mantini,  Dante and Coxon,  James P. and Swinnen,  Stephan P.},
  year = {2022},
  month = may,
  pages = {104338}
}

@article{Zhu2025,
  title = {In-phase and anti-phase dual-site beta tACS differentially influence functional connectivity and motor inhibition},
  url = {http://dx.doi.org/10.1101/2025.04.28.651005},
  DOI = {10.1101/2025.04.28.651005},
  publisher = {Cold Spring Harbor Laboratory},
  author = {Zhu,  Tingting and Sack,  Alexander T. and Leunissen,  Inge},
  year = {2025},
  month = may
}

@article{Schroll2018,
  title = {Reinforcement magnitudes modulate subthalamic beta band activity in patients with Parkinson’s disease},
  volume = {8},
  ISSN = {2045-2322},
  url = {http://dx.doi.org/10.1038/s41598-018-26887-3},
  DOI = {10.1038/s41598-018-26887-3},
  number = {1},
  journal = {Scientific Reports},
  publisher = {Springer Science and Business Media LLC},
  author = {Schroll,  Henning and Horn,  Andreas and Runge,  Joachim and Lipp,  Axel and Schneider,  Gerd-Helge and Krauss,  Joachim K. and Hamker,  Fred H. and K\"{u}hn,  Andrea A.},
  year = {2018},
  month = jun 
}

@article{Limongi2017,
  title = {Successful and unsuccessful response inhibitions differentially affect the effective connectivity between insular,  presupplementary-motor,  and striatal areas.},
  volume = {131},
  ISSN = {0735-7044},
  url = {http://dx.doi.org/10.1037/bne0000175},
  DOI = {10.1037/bne0000175},
  number = {1},
  journal = {Behavioral Neuroscience},
  publisher = {American Psychological Association (APA)},
  author = {Limongi,  Roberto and Pérez,  Francisco J.},
  year = {2017},
  pages = {20–32}
}

@article{Krause2014,
  title = {Not all brains are created equal: the relevance of individual differences in responsiveness to transcranial electrical stimulation},
  volume = {8},
  ISSN = {1662-5137},
  url = {http://dx.doi.org/10.3389/fnsys.2014.00025},
  DOI = {10.3389/fnsys.2014.00025},
  journal = {Frontiers in Systems Neuroscience},
  publisher = {Frontiers Media SA},
  author = {Krause,  Beatrix and Cohen Kadosh,  Roi},
  year = {2014}
}

@article{Alagapan2016,
  title = {Modulation of Cortical Oscillations by Low-Frequency Direct Cortical Stimulation Is State-Dependent},
  volume = {14},
  ISSN = {1545-7885},
  url = {http://dx.doi.org/10.1371/journal.pbio.1002424},
  DOI = {10.1371/journal.pbio.1002424},
  number = {3},
  journal = {PLOS Biology},
  publisher = {Public Library of Science (PLoS)},
  author = {Alagapan,  Sankaraleengam and Schmidt,  Stephen L. and Lefebvre,  Jérémie and Hadar,  Eldad and Shin,  Hae Won and Fr\"{o}hlich,  Flavio},
  editor = {Jensen,  Ole},
  year = {2016},
  month = mar,
  pages = {e1002424}
}

@article{Isherwood2025,
  title = {Multi-study fMRI outlooks on subcortical BOLD responses in the stop-signal paradigm},
  url = {http://dx.doi.org/10.7554/eLife.88652.3},
  DOI = {10.7554/elife.88652.3},
  publisher = {eLife Sciences Publications,  Ltd},
  author = {Isherwood,  Scott JS and Kemp,  Sarah A and Miletić,  Steven and Stevenson,  Niek and Bazin,  Pierre-Louis and Forstmann,  Birte U},
  year = {2025},
  month = jan 
}
