## Supplementary figures and images for "tACS induces state-dependent shifts in inhibitory phase preference consistent with phase precession"

### bioRxiv_logo.png

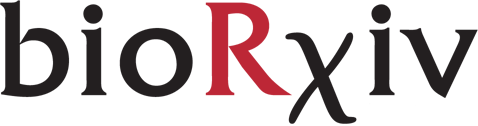

### fig1.jpg

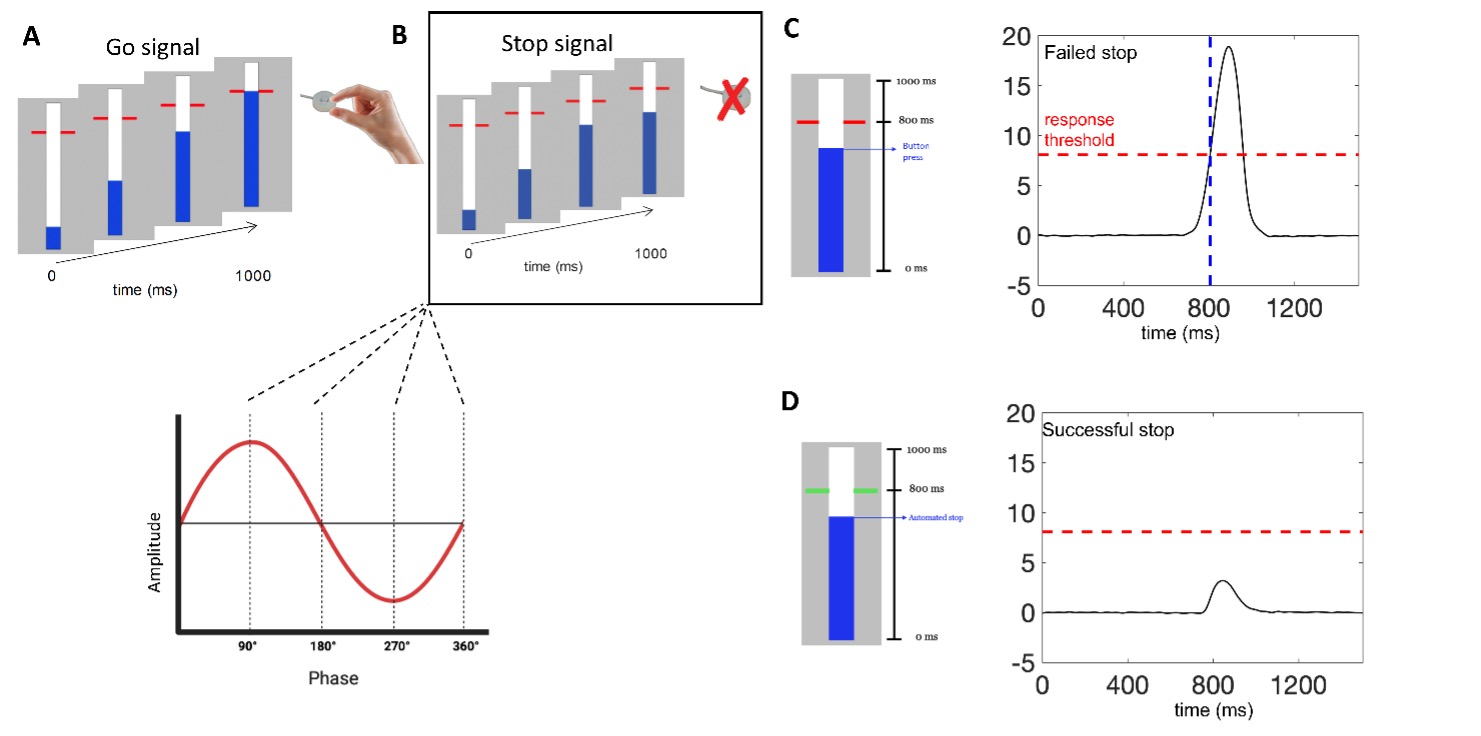

### fig2.jpg

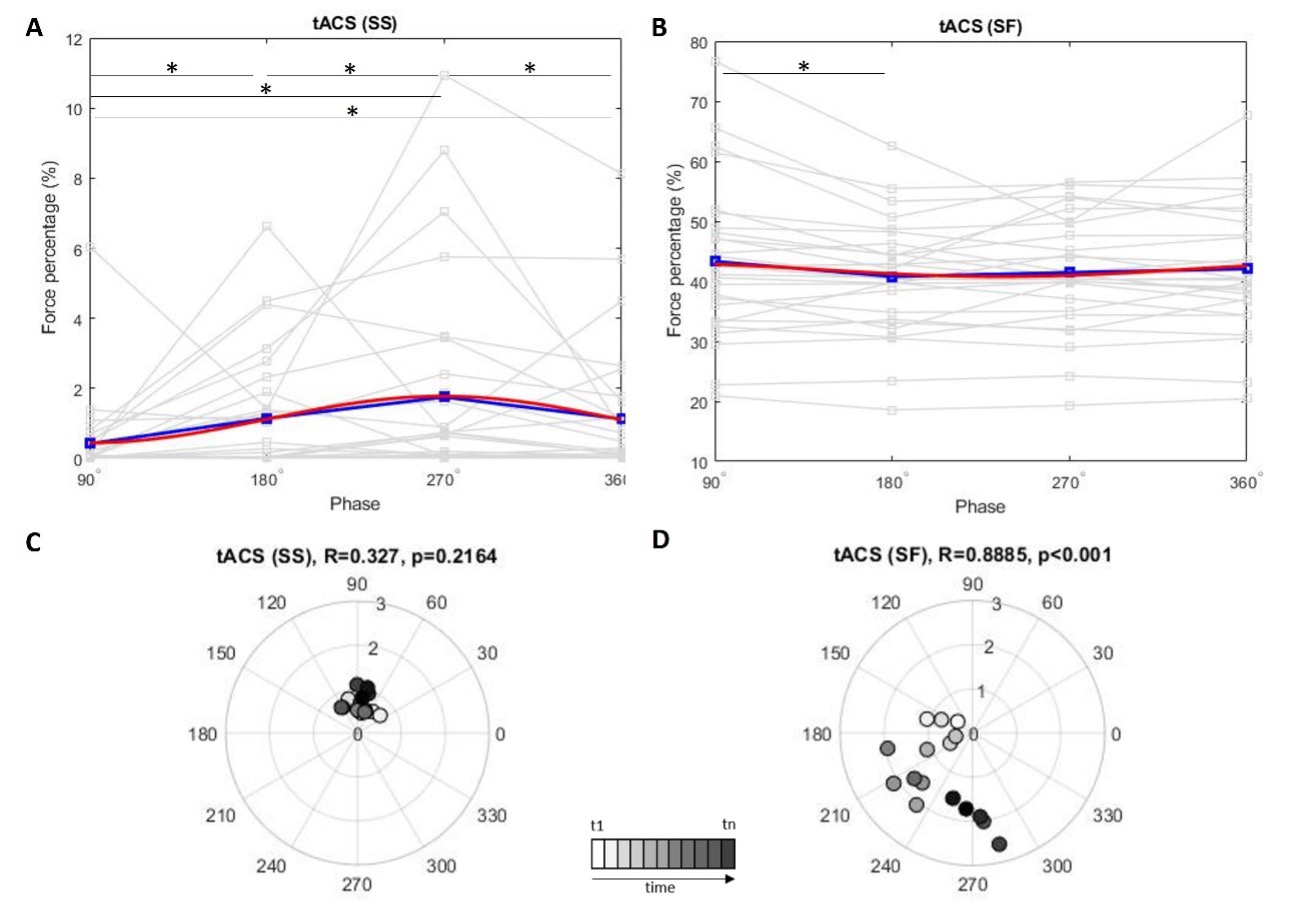

### fig3.jpg

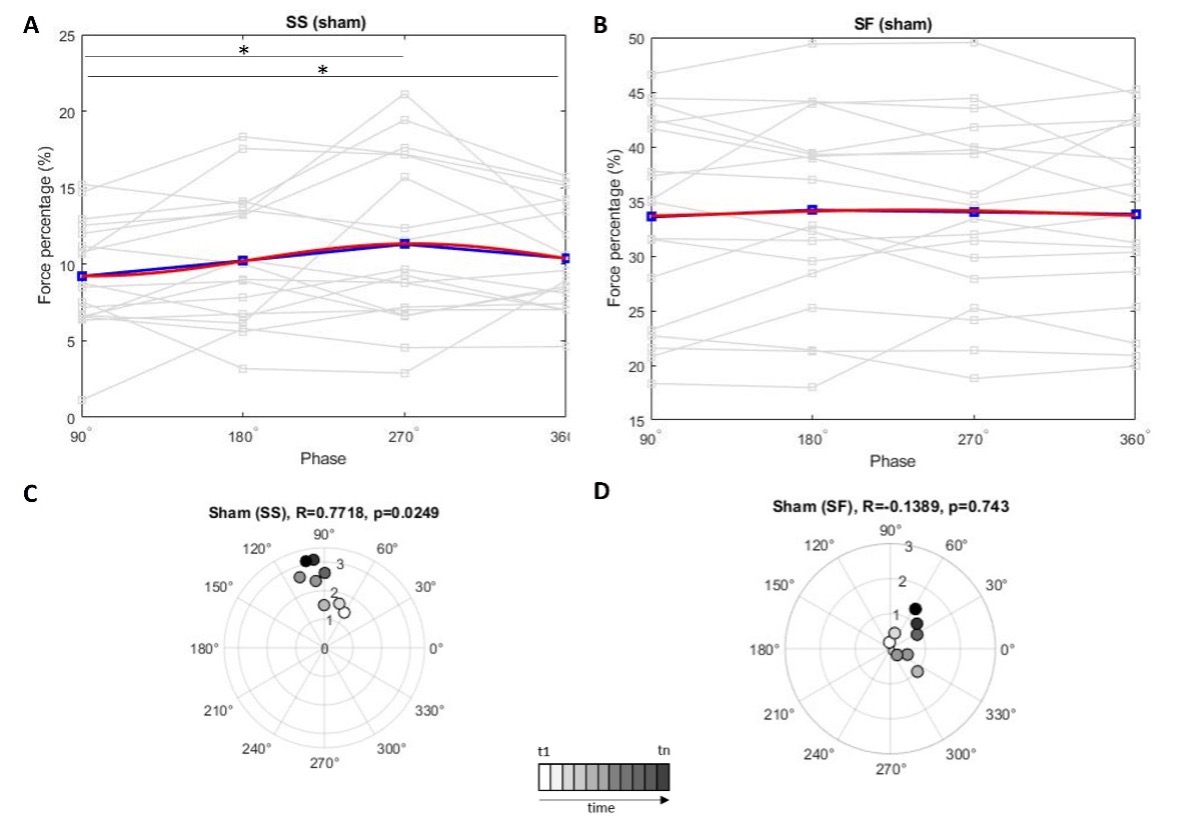

### figs1.jpg

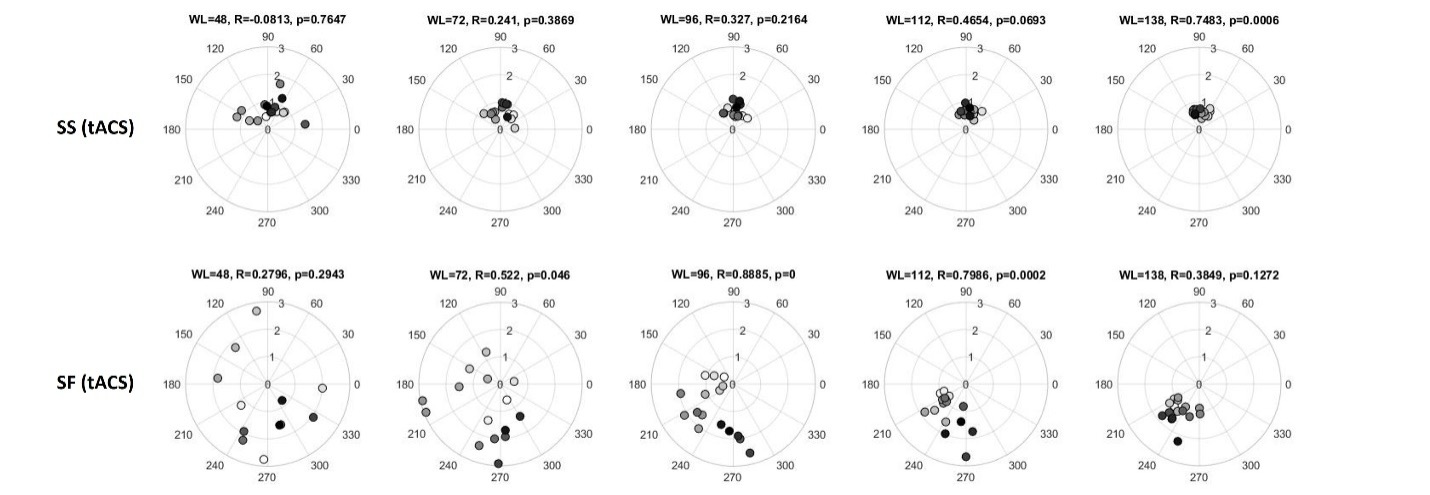

### figs2.jpg

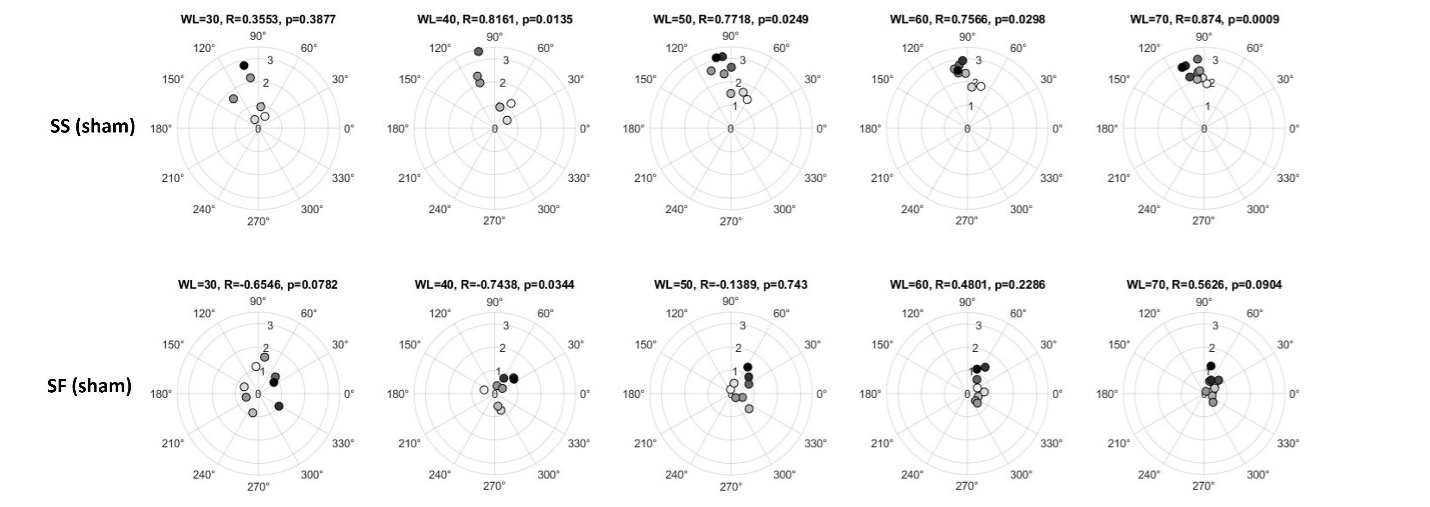

### figs3.jpg

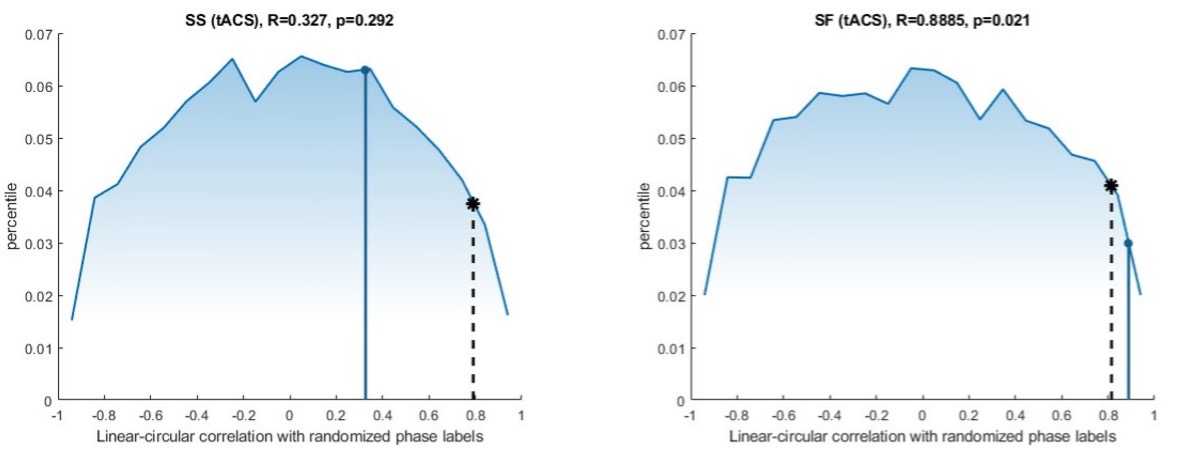

### figs4.jpg

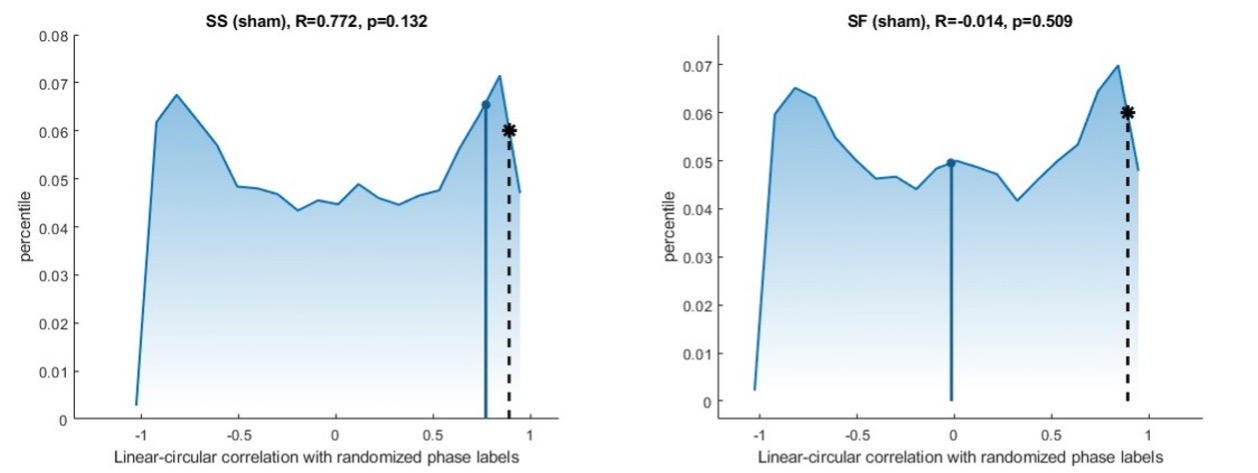
